# Bees attend primarily to costs, not benefits, to avoid exploitation by floral mimics

**DOI:** 10.1101/2025.11.20.689644

**Authors:** Annaliese N. Novinger, Claire T. Hemingway, Jenny K. Burrow, Charlotte C. Davis, Avery L. Russell

**Author notes:** Corresponding author: Department of Biology, 910 S John Q Hammons Pkwy, Temple Hall, Missouri State University, Springfield, MO 65897, USA. A.N.N. and C.T.H. are co-first authors. AUTHOR CONTRIBUTIONS: ANN and ALR conceived and designed the experiments. ANN, JKB, CCD, and ALR performed the experiments and collected the data. ANN and ALR analyzed the data. ANN made the illustrations in Figure 1. ANN, CTH, and ALR wrote the original draft of the manuscript; other authors provided editorial advice.

## Abstract

Animals are expected to weigh costs and benefits when making foraging decisions. Flowering plants often offer floral food rewards to pollinators, but also frequently deceive pollinators into visiting without providing rewards. Although floral mimicry, in which rewardless flowers mimic rewarding (model) flowers is common, it remains unclear whether pollinators evaluate the relative costs and benefits of visiting each. Using a signal detection theory (SDT) framework, we investigated how bumble bee decisions respond to experimental manipulations of costs and benefits within an intersexual floral mimicry system in which unrewarding female flowers imperfectly mimic pollen-rewarding male flowers (models). Using a fully factorial design, we increased the cost of visiting mimics (by adding quinine), increased the benefit of visiting models (by adding pollen), increased both simultaneously, and made no changes (control). As predicted by SDT, increasing mimic costs led to a conservative bias: bees made more correct rejections, but fewer correct detections. Contradicting SDT predictions, enhancing model reward did not elicit a liberal bias and decision performance remained unchanged. Our findings suggest that bees prioritize avoiding costly errors under uncertainty, even when this limits potential gains. Such conservative foraging decisions may help stabilize deceptive systems by allowing mimics to persist despite moderate costs.

## INTRODUCTION

Animals often make decisions under uncertainty, continually updating information about their environment to reduce ambiguity and optimize choice behavior (Blumstein & Bouskila, 1996; Real, 1992; Stephens, 2008). For pollinators such as bees, this ability is critical. During a single day, a bee may visit thousands of flowers across multiple plant species that vary in their floral traits and reward payoffs (Chittka et al., 1999; Heinrich, 1976, 1979). To maximize energetic return, pollinators must rapidly and accurately assess complex and variable floral signals – yet these signals are not always reliable. Because producing floral rewards is energetically costly (Nicholls & Hempel De Ibarra, 2017; Pyke & Ren, 2023; Southwick, 1984), plants may use floral signals to attract visitors without offering rewards. This occurs in Batesian floral mimicry, in which a rewardless plant species (mimic) produces deceptive signals that overlap in sensory space with those of a rewarding species (model), in one or more sensory modalities (Dafni, 1984; Johnson & Schiestl, 2016; Pannell & Farmer, 2016). This form of deception is widespread, having evolved repeatedly across angiosperms in more than 32 plant families and across 7,500 species worldwide (Johnson & Schiestl, 2016; Renner, 2006; Schiestl & Johnson, 2013). The success of this strategy depends on the perceptual and learning limitations of pollinators, which may fail to reliably discriminate mimic from model and thus visit both (Ferdy et al., 1998; Renner, 2006; Schiestl & Johnson, 2013; Vázquez & Barradas, 2017).

Signal detection theory (SDT) offers a powerful framework for understanding why pollinators sometimes visit rewardless flowers when faced with uncertainty (Lichtenberg et al., 2020). When pollinators encounter deceptive mimics, they face four possible decision outcomes (Figure 1): two correct decisions – visiting a model (‘hit’) and avoiding an unrewarding mimic (‘correct rejection’) – and two types of decision errors – visiting a rewardless flower (‘false alarm’) or failing to visit a rewarding one (‘miss’). SDT describes the tradeoff between these two types of decision errors (Green & Swets, 1966; Wiley, 2006). Because reducing one type of error increases the other, pollinators must adopt a decision criterion that balances the costs and benefits of these errors. While SDT has been widely used to explain the evolution and maintenance of floral mimicry, most studies have focused on the ability of pollinators to discriminate between model and mimics based on their perceptual similarity (e.g., Gumbert & Kunze, 2001; Leonard et al., 2011; Russell et al., 2021) and on the relative frequency of each within a given population (e.g., Finkbeiner et al., 2018; Lindström et al., 1997; Pfennig et al., 2001; Russell et al., 2020). Much less is known about how the costs and benefits of foraging outcomes shape pollinator decision criteria.

**Figure 1.**
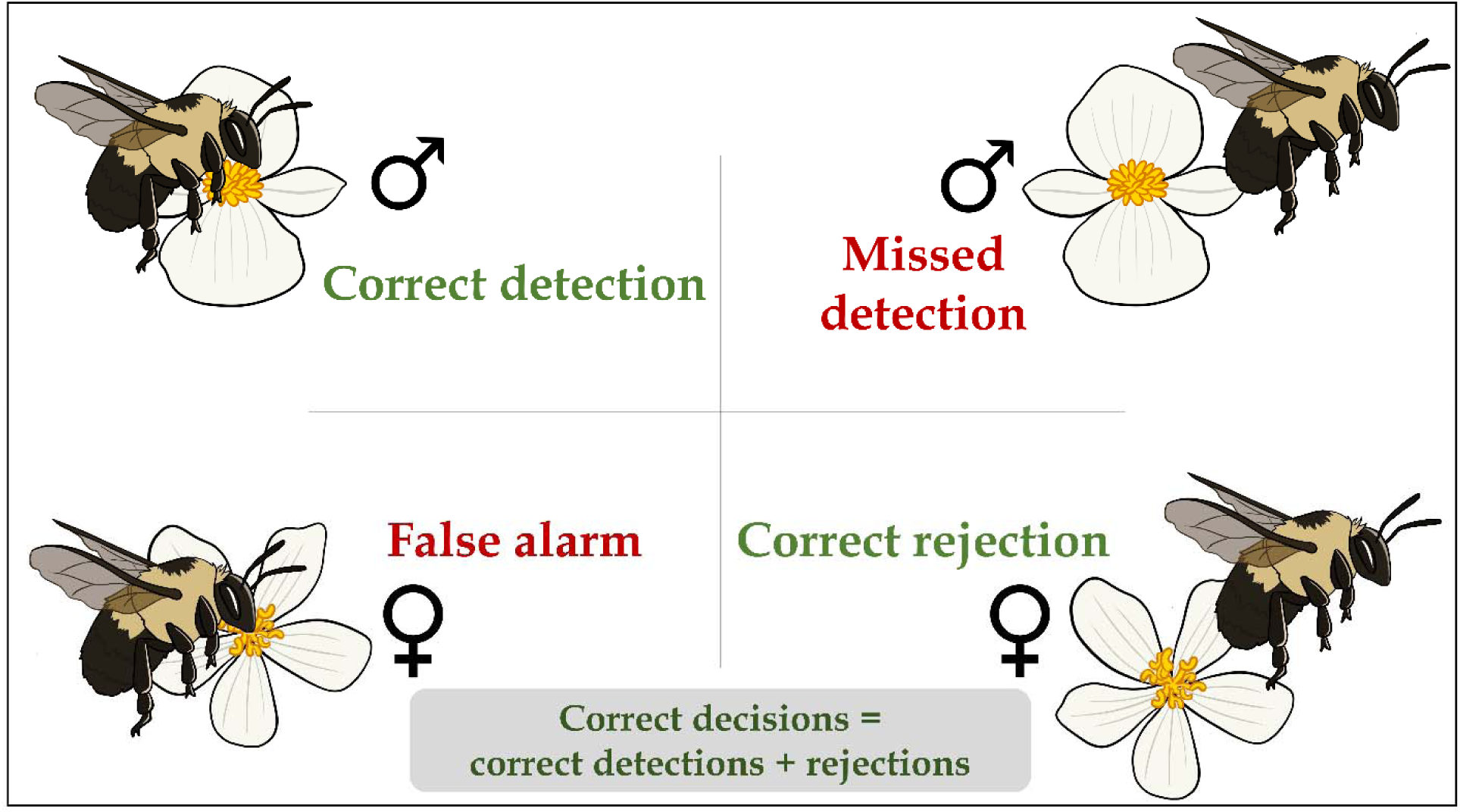
Sampling behavior of bees foraging on the intersexual floral mimic *Begonia odorata* can be divided into four categories in accordance with signal detection theory. **Correct detections** (‘hits’) involve the bee approaching and landing on models, whereas **missed detections** (‘misses’) involve the bee approaching, but not landing on models. **Correct rejections** involve the bee approaching, but not landing on mimics, whereas **false alarm** involve the bee approaching and landing on mimics. Thus, **correct decisions** involve the bee approaching and landing on models, and approaching, but not landing on mimics.

This is somewhat surprising, given a central prediction of SDT is that receivers should adjust their decision thresholds according to the relative costs and benefits of foraging on models and mimics (Abbott & Sherratt, 2013; Kikuchi & Sherratt, 2015; Sherratt & Peet-Paré, 2017). For pollinators, foraging carries clear costs, including energetic expenditure and time investment (Heyneman, 1983; Mayberry et al., 2024; Wolf et al., 1975), exposure to predators (Romero et al., 2011), and sometimes contact with toxic or otherwise aversive floral rewards (Barlow et al., 2017; Jones & Agrawal, 2016; Stevenson et al., 2017). The benefits of foraging are also variable, depending on the nutritional value, quantity, and temporal or spatial availability of rewards (Benadi et al., 2023; Hemingway, Leonard, et al., 2024; Venjakob et al., 2022). Although pollinators are sensitive to these costs and benefits while foraging (e.g., Hemingway et al., 2024; Richman et al., 2017), their influence on decision making in deceptive systems remains poorly understood (see de Jager & Ellis, 2014; J. Ferdy et al., 1998; Gaskett, 2011; Russell et al., 2020). SDT predicts plasticity in criteria: bees should be more cautious (avoid false alarms) when errors are costly, or more liberal (favor hits) when rewards are high. Moreover, when both the cost of visiting the mimic and the benefit of visiting the model are increased, the bias should reflect the combined influence of costs and benefits (Lynn & Barrett, 2014).

Here we used a powerful system to empirically test these predictions of SDT within the context of floral deception. *Begonia odorata* is a monoecious species bearing both male and female flowers on the same plant, and these flowers are visually similar, with showy white tepals presented together within inflorescences (Russell et al., 2020). This system exhibits intersexual floral mimicry, in which unrewarding female flowers mimic pollen-producing male flowers – a phenomenon known as pseudoanthery (Johnson & Schiestl, 2016). Pollinators, typically pollen-collecting bees, learn to associate male-phase traits (such as color, scent, and shape) with pollen rewards, creating a signal-reward mismatch that requires them to discriminate between visually similar morphs. Previous work has shown that naïve bees exhibit a sensory bias toward mimics but rapidly learn to avoid them, and that model-mimic frequency has little effect on discrimination performance (Russell et al., 2020). Subsequent experiments tested whether variation in flower size influenced the rate of discrimination learning and found that although bees quickly learned to distinguish between rewarding males and unrewarding females, this learning was not significantly affected by size variation (Russell et al., 2021). While clearly bees can readily learn to discriminate models from mimics, it remains unclear how discrimination is shaped by the relative rewards of models and the costs of visiting mimics. *Begonia odorata* therefore provides an ideal system for testing how variation in costs and benefits influences pollinator decision-making.

In a fully factorial experiment, we manipulated the rewards of male flowers (via addition of pollen) and the punishments of female flowers (via addition of aversive quinine powder) to vary the payoffs of hits and false alarms. We asked how these manipulations, and pollinator experience, affect bees’ decision biases and discrimination abilities. Consistent with SDT, we predicted a conservative bias in bee decision making when the cost of visiting mimics was increased and a liberal bias when the benefit of visiting models was increased, and that bias would reflect the sum of the cost plus the benefit when both costs and benefits of mimics and models were increased, respectively. As a result, we predicted that when mimic cost increased, bees would make relatively more correct rejections at the cost of making fewer correct detections (‘hits’) and would be less likely to buzz flowers, an energetically expensive attempt to extract pollen (Rossi et al., 2025). Likewise, we predicted that when model benefit increased, bees would make relatively more correct detections, at the cost of fewer correct rejections, and would be more likely to buzz flowers. Finally, as naïve pollinators are initially unfamiliar with the phenotypic distribution and costs and benefits of visiting models and mimics, we expected bees’ responses to models and mimics to increasingly fit these predictions of SDT as foraging bees gained experience (e.g., (Russell et al., 2020, 2021).

## METHODS

### Test subjects

We maintained 3 colonies (Plant Products: Biobest Group, Canton, MI, U.S.A.) of the common eastern bumble bee *Bombus impatiens* following Russell et al., 2020. In brief, we allowed colonies to forage freely on 2 M sucrose solution and pulverized honeybee-collected pollen (Koppert Biological Systems) from artificial feeders within enclosed foraging arenas (length, width, height: 82 × 60 × 60 cm) set to a 14 h : 10 h light : dark cycle.

We used fresh male and female flowers with mature anthers and styles, respectively, from 10 simultaneously monoecious *Begonia odorata* plants raised in a university greenhouse with supplemental halogen lights to extend day length to a 14 : 10 h cycle and with fertilizer applications every other week (Plant Tone, NPK 5 : 3 : 3). While female *B. odorata* flowers are rewardless and do not produce pollen or nectar, male *B. odorata* flowers offer pollen, their only reward to their primary pollinators, bees; bumble bees are among the bee genera that visit closely related *Begonia* species (De Avila et al., 2017; Pemberton & Wheeler, 2006; Schemske et al., 1996; Wyatt & Sazima, 2011). Female *B. odorata* flowers closely resemble male flowers in bumble bee color vision (Russell et al., 2020, 2021) and scent (unpublished data); both flower sexes have creamy white dissected petals, and the female flower’s yellow and highly divided styles closely resemble the male flower’s numerous yellow stamens.

### Experimental overview

We tested how the benefits of visiting models (male flowers) and the costs of visiting mimics (female flowers) influenced learning in initially flower-naïve bees (*N* = 97). We examined how bees learned to sample between models and mimics by analyzing three primary components of their sampling behavior during visits to arrays of 20 flowers: approaches, landings without buzzing, and landing with buzzing (“buzzes”). An approach was defined as a bee in flight greatly reducing its velocity while facing a flower within 3 cm. All landings (defined as 3 or more legs in contact with the flower) were preceded by an approach. Landings were considered correct detections when directed at models and false alarms when directed at mimics (Figure 1). Not all approaches led to landings; these cases represented missed detections for models and correct rejections for mimics (Figure 1). Landings without buzzing on male flowers typically involved the bee collecting pollen via a behavior termed scrabbling; see Russell et al., 2017 for a description. In contrast, bees never scrabbled on female flowers. Buzzes, which indicated an attempt at extracting pollen whether or not it was available, were identified by their distinctive sound using Zoom H2 Handy Recorders (ZOOM Corporation) to amplify and verify buzzes while trials were taking place and occurred only after a bee had landed (see Russell et al. 2016a). Buzzing a male flower represented a correct response, whereas buzzing a female flower represented an incorrect response.

We assigned flower-naïve bees approximately evenly across five treatments: an unmodified control treatment, a modified control treatment (‘control treatment’), an increased model benefit treatment (‘benefit treatment’), an increased mimic cost treatment (‘cost treatment’), and an increased model benefit and mimic cost treatment (‘both treatment’). In the unmodified control treatment, both male flowers (which offer on average 0.18 mg of pollen) and female flowers were presented to bees without modification. In all other treatments, anthers of male flowers were supplemented with pollen. Because collecting pollen from male *Begonia* flowers to supplement other flowers was extremely time intensive, we used cherry pollen for all supplementation (*Prunus avium*; Pollen Collection and Sales, Lemon Cove, California, USA), which bumble bees readily collect (e.g., Russell et al., 2017). In the ‘control treatment’, male flowers were each supplemented with 0.9 mg pollen (to control for the effect of adding a novel pollen type across treatments) and female flowers were unmodified. To increase the benefit to bees when visiting models, in the ‘benefit treatment’, male flowers received double the pollen supplement (1.8 mg pollen added to the anthers of each flower), while female flowers were unmodified. To increase the cost to bees when visiting mimics, in the ‘cost treatment’, the styles of female flowers were supplemented with 0.5 mg of powdered quinine (99% anhydrous, ACROS Organics, Fisher Scientific, Hampton, NH, USA), a stimulus that bees perceive as aversive (Muth et al., 2016), while male flowers received the base supplement (0.9 mg pollen). In the ‘both treatment’, male flowers received the double pollen supplement (1.8 mg), while female flowers received the quinine supplement. Across all treatments, 20 flowers were arranged in a 5 x 4 Cartesian grid, with flowers spaced 7 cm apart. Male and female flowers were alternated by position and presented in equal numbers. To prevent desiccation, freshly cut live flowers were placed into custom water tubes (Russell et al., 2017). We systematically alternated trials among treatments to control for potential effects of time and day.

To initiate a behavioral trial, flowers were set up and a single flower-naïve worker bee was gently captured from the foraging arena using a 40 dram vial (Bioquip) and immediately released in the center of the test arena following (Russell et al., 2017). We terminated the trial after 60 visits (or earlier if the bee stopped visiting flowers for 5 minutes) to avoid bees depleting models of pollen rewards (mean 47 visits; range 4-60 visits). To terminate the trial, we captured the bee in a 40-dram vial and euthanized it. For each trial we used a new flower-naïve bee and new freshly clipped flowers; we tested bees individually and never reused flowers or bees across trials. The test arena was cleaned after each trial. For analyses, we excluded one bee with fewer than five visits and no flower landings.

### Data analyses

All data were analyzed using R v.4.4.0 (Developmental Core Team R, 2024). To analyze how experience and the costs of visiting mimics and benefits of visiting models affected sampling behavior, we fit generalized linear mixed models (GLMMs) with a binomial distribution using the glmmTMB() function (glmmTMB package; Brooks, 2025), specifying type II Wald chi-squared (χ2)-tests via the Anova() function (car package; Fox et al., 2023). We checked model assumptions for all models (DHARMa package; Hartig, 2024).

For the first set of GLMMs, the response variable was sampling behavior (either ‘correct decisions’ vs ‘incorrect decision’, ‘correct detection’ vs ‘missed detection’, ‘correct rejection’ vs ‘false alarm’, ‘buzz given landing’ vs ‘land without buzzing’, or ‘landing on male’ vs ‘landing on female’) and the explanatory variables were ‘treatment’ (modified control (‘Control’), increased model benefit (‘Benefit’), increased mimic cost (‘Cost’), and increased mimic cost and model benefit (‘Both’) and experience (‘visit number’). We included ‘bee’ as a random factor, with ‘visit number’ as repeated measures within bee, within colony (except for two models, in which bee within colony would not converge; we therefore excluded the colony random effect for these two models). In cases of significant effects, we ran Tukey’s post hoc test using the emmeans() function (emmeans package; Length, 2025) to determine which pairs were significant. For the second set of GLMMs, in which we examined how pollen reward supplementation specifically affected sampling behavior, the explanatory variable ‘treatment’ was either the unmodified control or the modified control.

To determine whether pollen or quinine supplementation affected flower-naïve bees’ initial landing preference for models (male flowers) versus mimics (female flowers), we used G-tests (DescTools package; Signorell et al., 2025) to examine choice of first flower landing. There were no significant differences in model versus mimic preference for any treatment (modified control: *G* = 0.430, *N* = 21, *P* = 0.512; increased model benefit: *G* = 1.33, *P* = 0.249, *N* = 19; increased mimic cost: *G* = 0.00, *P* = 1.000, *N* = 20; increased mimic cost and model benefit: *G* = 0.20, *P* = 0.655, *N* = 21).

## RESULTS

### Correct decisions and detections are affected primarily by mimic cost

Flower-naïve bees overall learned to make proportionally more correct decisions (combining correctly rejecting mimics and correctly detecting models) with experience, but this pattern depended on treatment: increasing the cost of mimics modestly reduced the proportion of correct decisions with experience (Figure 2a; GLMM: treatment effect: χ^2^3 = 19.71, *P* < 0.0002; experience effect: χ^2^1 = 6.38, *P* < 0.012; treatment × experience effect: χ^2^3 = 10.06, *P* < 0.019). Bees made proportionally fewer correct detections with experience and the rate of correct detections decreased much more quickly when mimics were relatively more costly to visit (Figure 2b; GLMM: treatment effect: χ^2^3 = 45.79, *P* < 0.0001; experience effect: χ^2^1 = 96.97, P < 0.0001; treatment × experience effect: χ^2^3 = 8.57, *P* < 0.036). Bees made proportionally more correct rejections with experience, and this pattern did not depend on the cost of mimics and benefit of models (Figure 2c; GLMMs: treatment effect: χ^2^3 = 6.94, *P* = 0.074; experience effect: χ^2^1 =130.07, *P* < 0.0001; treatment × experience effect: χ^2^3 = 0.67, *P* = 0.880).

### Decision to buzz is affected primarily by mimic cost

Bees did not alter how often they buzzed models with experience, but overall buzzed models significantly less when the cost of visiting mimics was increased (Figure 3a; GLMM: treatment effect: χ^2^_3_ = 66.49, *P* < 0.0001; experience effect: χ^2^1 = 1.56, *P* = 0.213; treatment × experience effect: χ^2^3 = 3.57, *P* = 0.313). Bees altered how often they buzzed mimics with experience, but this pattern depended on the cost of visiting mimics (Figure 3b). When the cost of mimics was unmodified, bees initially buzzed frequently, but less so with experience; when the cost of mimics was increased, bees initially rarely buzzed, but increased buzzing somewhat with experience (Figure 3b; GLMM: treatment effect:χ^2^3 = 43.56, *P* < 0.0001; experience effect: χ^2^1 = 20.07, *P* < 0.0001; treatment × experience effect: χ^2^3 = 31.29, *P* < 0.0001). Finally, the proportion of landings on models changed with experience, but this relationship was complex (Figure 3c). Landings on models increased the most when only the benefit of visiting models was increased; landings on models decreased the most when costs of mimics and benefits of models were increased simultaneously (Figure 3c; GLMM: treatment effect: χ^2^3 = 4.94, *P* = 0.177; experience effect: χ^2^1 = 2.96, *P* < 0.086; treatment × experience effect: χ^2^3 = 8.41, *P* < 0.039).

**Figure 2.**
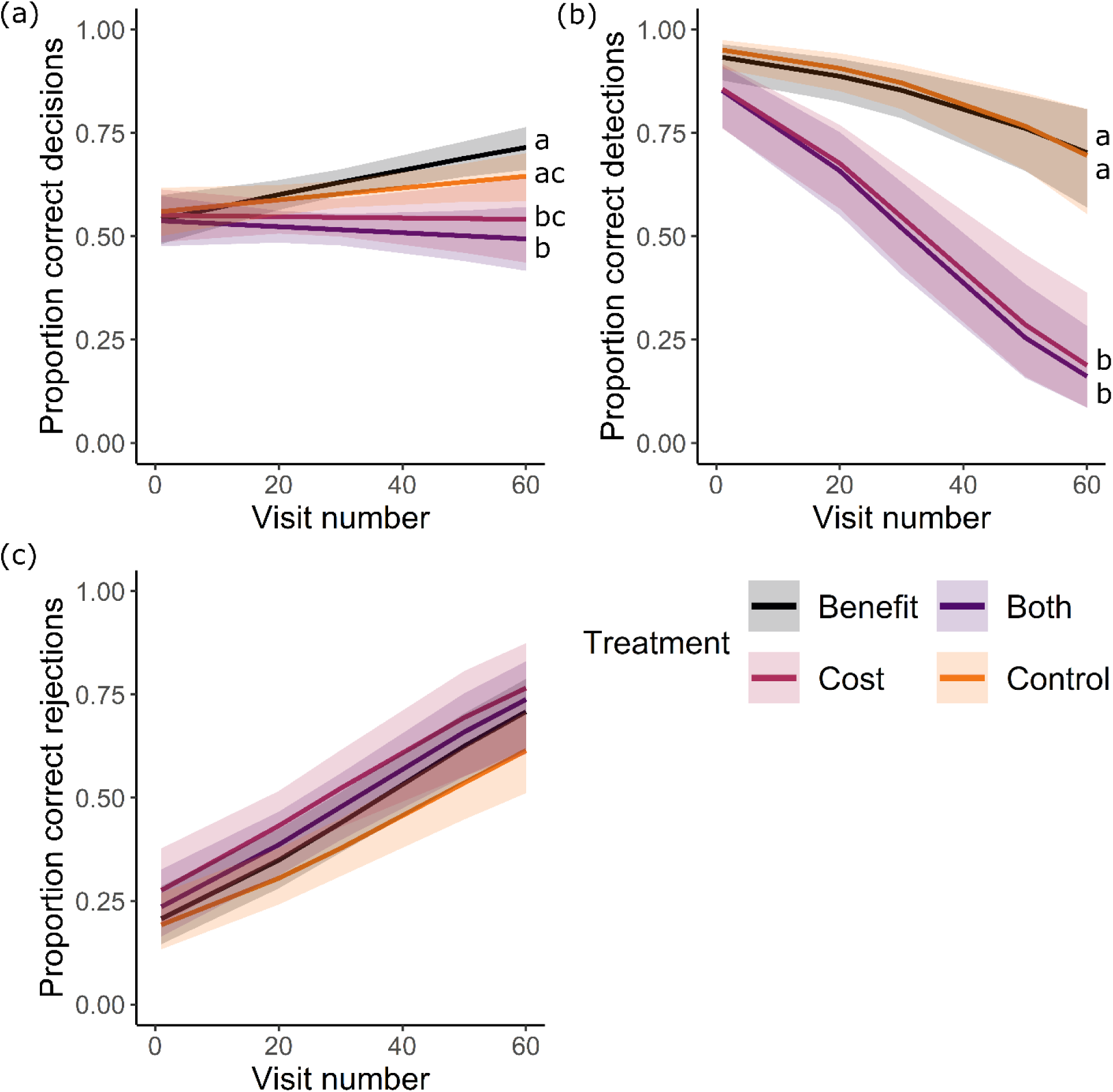
Sampling behavior of initially naïve bees foraging in treatments (different colored lines) that differed in terms of the relative costs and benefits of mimics and models, respectively. The mean proportion of (a) correct decisions, (b) correct detections, and (c) correct rejections made by bees over the course of up to 60 visits. *N* = 21, 19, 20 and 21 bees in the modified control (‘Control’), increased model benefit (‘Benefit’), increased mimic cost (‘Cost’), and increased mimic cost and model benefit (‘Both’) treatments, respectively. Plotted lines indicate estimated means and shaded regions indicate 95% confidence intervals. Different letters next to plotted lines indicate significant differences among treatments at *P* < 0.05 according to Tukey’ post hoc tests.

**Figure 3.**
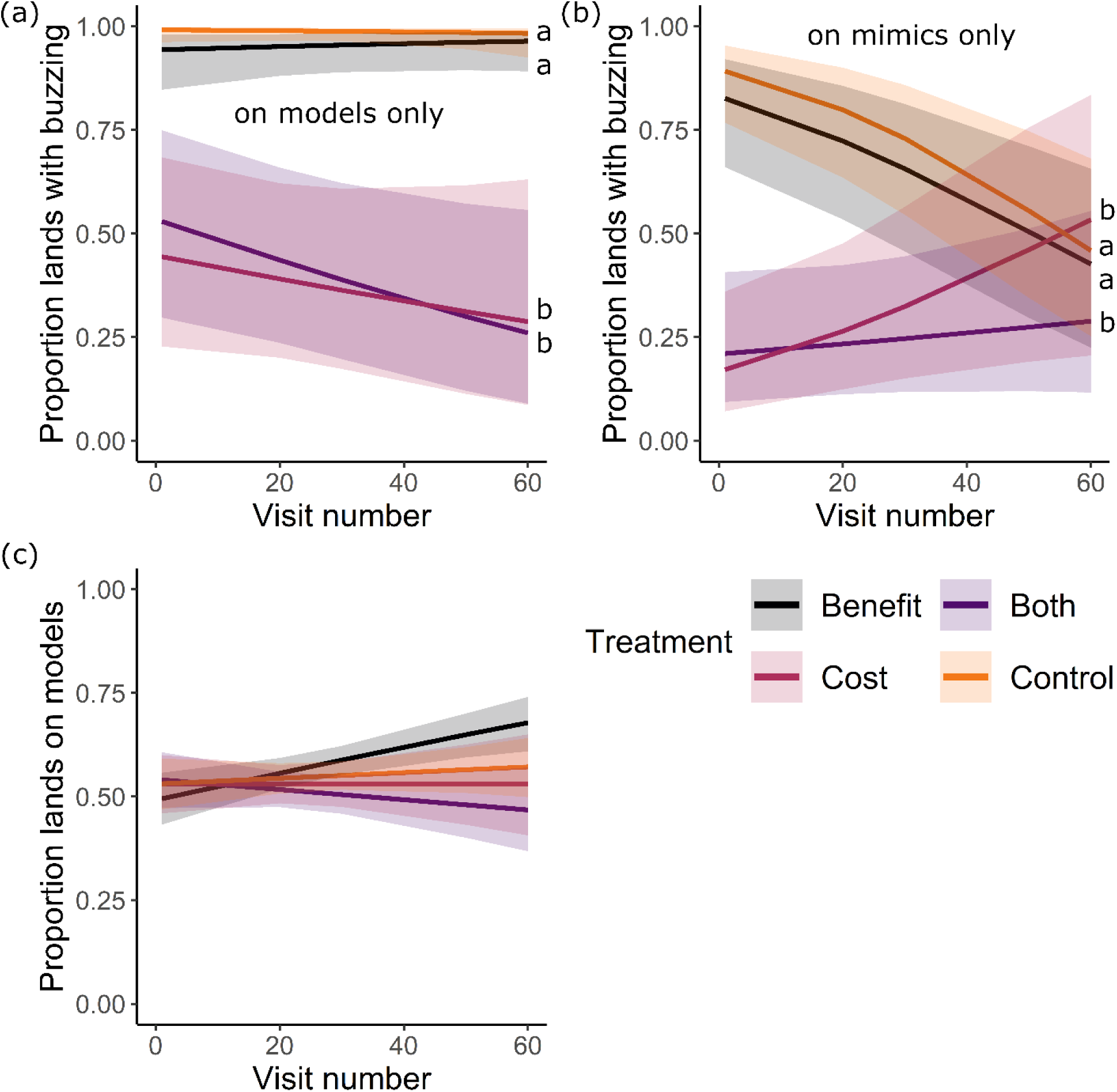
Landing behavior of initially naïve bees (same dataset as in Figure 2, analyzed for different sampling behavior) foraging in treatments (different colored lines) that differed in term of the relative costs and benefits of mimics and models, respectively. Mean proportion of lands made on (a) models during which the bee buzzed, (b) mimics during which the bee buzzed, and on (a) models (versus mimics), over the course of up to 60 visits. *N* = 21, 19, 20 and 21 bees in the modified control (‘Control’), increased model benefit (‘Benefit’), increased mimic cost (‘Cost’), and increased mimic cost and model benefit (‘Both’) treatments, respectively. Plotted lines indicate estimated means and shaded regions indicate 95% confidence intervals. Different letters next to plotted lines indicate significant differences among treatments at *P* < 0.05 according to Tukey’s post hoc tests.

### Decision making is modestly improved when models are pollen supplemented

Bees made proportionally more correct decisions with experience but made consistently more correct decisions (15% more on average) when models were supplemented with pollen (Figure 4a; GLMM: treatment effect: χ^2^1 = 11.55, *P* < 0.0007; experience effect: χ^2^1 = 3.96, *P* < 0.047; treatment × experience effect: χ^2^1 = 0.13, *P* = 0.719). Bees decreased their proportion of correct detections with experience but made consistently more correct detections (7.5% more on average, though this difference was not statistically significant) when models were supplemented with pollen (Figure 4b; GLMM: treatment effect: χ^2^1 = 2.57, *P* = 0.110; experience effect: χ^2^1 = 18.80, *P* < 0.0001; treatment × experience effect: χ^2^1 = 3.55, *P* = 0.060). Bees increased their proportion of correct rejections with experience but made consistently more correct rejections (30% more on average) when models were supplemented with pollen (Figure 4c; GLMM: treatment effect: χ^2^1 = 6.40, *P* < 0.0099; experience effect: χ^2^1 = 43.05, *P* < 0.0001; treatment × experience effect: χ^2^1 = 2.58, *P* = 0.108). Bees in both treatments slightly improved their proportions of landings on models with experience, with bees making consistently more landings on models (13% more on average) when supplemented with pollen (Figure 4d; GLMM: treatment effect: χ^2^1 = 5.82, *P* < 0.016; experience effect: χ^2^1 = 0.60, *P* = 0.438; treatment × experience effect: χ^2^1 = 0.044, *P* = 0.834).

## DISCUSSION

When encountering floral mimicry, pollinators must continually discriminate between rewarding and unrewarding flowers, often under conditions of incomplete information and uncertainty (Johnson & Schiestl, 2016). Using a framework derived from signal detection theory (SDT), we asked how increasing decision costs and benefits influenced bees’ decision criteria. We found that bees adjusted their decision thresholds in response to increased costs, but not to increased benefits. When the cost of encountering a mimic was elevated by adding quinine, bees exhibited a conservative bias, producing more correct rejections and fewer correct detections; bees also made far fewer costly attempts to collect pollen (Rossi et al., 2025) on models and mimics. However, increasing rewards associated with male flowers by providing extra pollen did not elicit a corresponding liberal shift. This asymmetry suggests that bees are more responsive to potential costs than to benefits. This pattern only partially aligns with the predictions of signal detection theory, which predicts that receivers should adjust their decision thresholds according to the relative costs and benefits of foraging on models and mimics (Abbott & Sherratt, 2013; Kikuchi & Sherratt, 2015; Sherratt & Peet-Paré, 2017).

**Figure 4.**
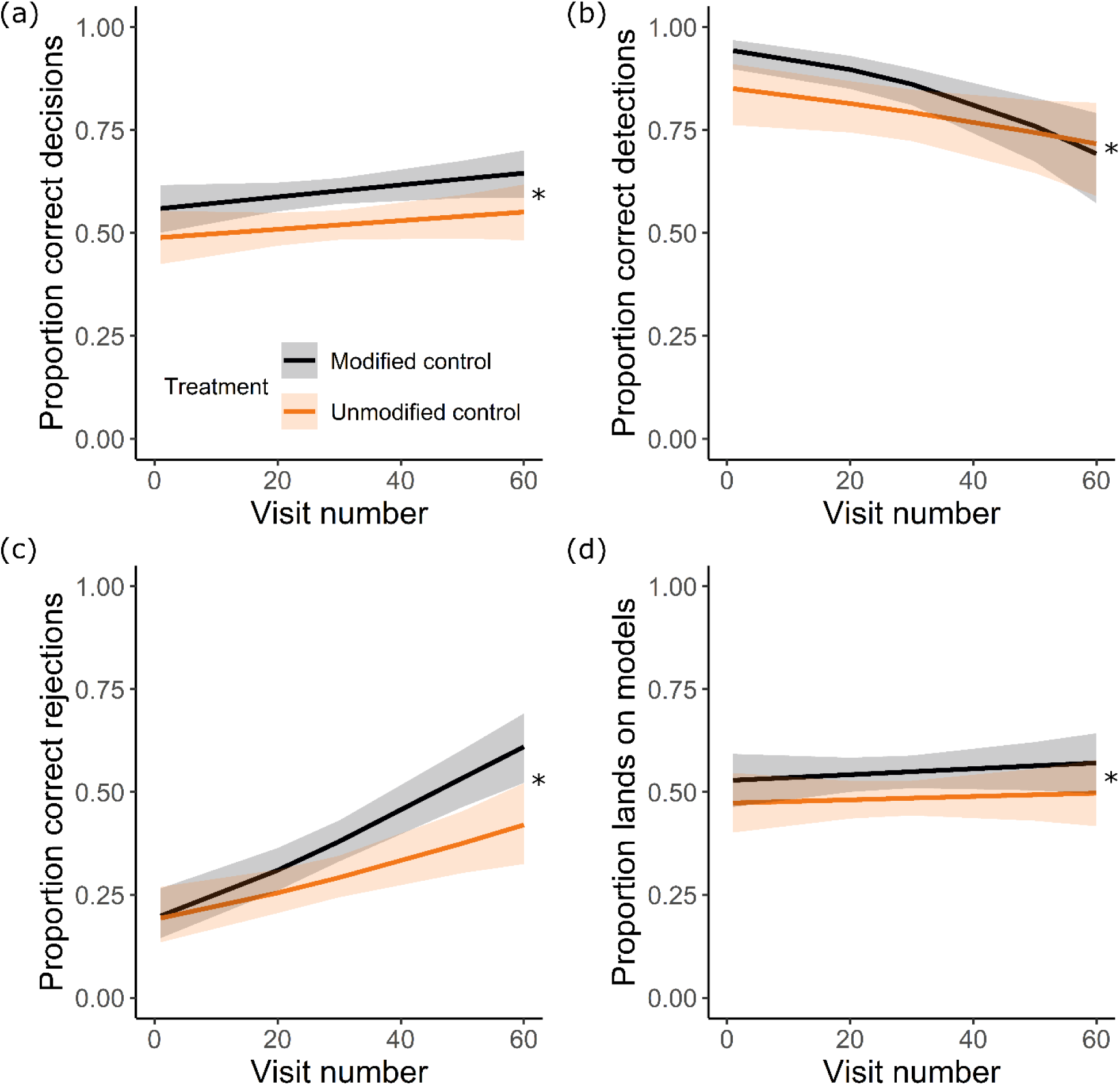
Sampling and landing behavior of initially naïve bees foraging in treatments differing in terms of whether we supplemented pollen to models (partially different dataset from Figures 2 and 3, analyzed for same behaviors). Mean proportion of (a) correct decisions, (b) correct detections, (c) correct rejections, and (d) lands made on models (versus mimics), over the cours of up to 60 visits. *N* = 21 and 16 bees in the pollen supplemented (‘Modified Control’) and natural (‘Unmodified Control’) treatments, respectively. Plotted lines indicate estimated means and shaded regions indicate 95% confidence intervals. Asterisks indicate significant differences among treatments at *P* < 0.05 according to Tukey’s post hoc tests.

More broadly, these findings align with published evidence that bees tend to minimize costly errors, even when this strategy limits potential gains. In many contexts, when foraging decisions are made under uncertainty, individuals must balance the expected payoff of each option with the variability in outcomes – commonly referred to as risk-sensitivity (Kacelnik & Bateson, 1996; McNamara & Houston, 1992). Such models predict that animals evaluate both the mean and variability of rewards, with risk-averse individuals favoring consistent outcomes and risk-prone individuals favoring variability when average gains are equivalent. Many animals exhibit risk-prone behavior when facing delays but are generally risk-averse or risk-neutral when outcomes differ in magnitude or probability, including cases where rewards may be absent (Kacelnik & Bateson, 1996). Bumble bees exhibit risk aversion when nectar volumes vary (Harder & Real, 1987), but individuals may shift toward being risk-prone when the colony’s energetic reserves are low (Cartar & Dill, 1990). Similarly, honey bees become more risk-averse when discrimination among rewards becomes more difficult, suggesting that uncertainty may amplify avoidance of costly mistakes (Shafir et al., 2008). Considered together with our findings, bees may generally prioritize minimizing costly errors, such as visiting unrewarding or punishing mimics, over maximizing potential gains, potentially reflecting a broader conservative bias in their decision-making under uncertainty.

Here, we made female (mimic) flowers more aversive by adding quinine. It remains unclear whether similar patterns of discrimination would emerge if the costs associated with floral deception were altered in different ways; however, we suspect that such manipulations could produce even stronger effects. In natural foraging environments, pollinators encounter a range of costs associated with decision errors, including energetic, time, and missed opportunities, all of which likely have fitness consequences (de Jager & Ellis, 2014; Nakazawa et al., 2024; Raine & Chittka, 2008; Zurbuchen et al., 2010). Pollinators are also susceptible to predator attacks while foraging, which can have serious fitness consequences (Romero et al., 2011). Increasing any of these costs may shift pollinator decision-making toward a more conservative strategy in which they avoid potential mimics, even at the expense of occasionally rejecting rewarding models. Under conditions where costs become sufficiently high, such as when deceptive flowers are abundant (Duffy & Johnson, 2017; Ferdy et al., 1999) or rewards are consistently poor, pollinators may even stop visiting either flower type altogether.

Conversely, what if the benefits were enhanced? For pollinators, the primary rewards offered by flowers are nectar and pollen, and increases in either their quantity or quality generally promote higher visitation rates and greater resource collection (Cartar, 2004; Hemingway et al., 2024; Nachev et al., 2017; Robertson et al., 1999; Ruedenauer et al., 2016). Here, we increased the reward value of male (model) flowers by supplementing them with pollen. It seems unlikely that adding even more pollen would increase the perceived benefits associated with male flowers, as bees did not deplete all pollen in most trials. Pollen supplementation relative to flowers in their natural state however did increase bee performance overall, suggesting that pollen odor acts as an honest cue of reward presence, potentially explaining why male *Begonia* offer so little pollen per flower. Changes in reward quantity may be perceived as less beneficial than changes in quality (see Cnaani et al., 2006), and these patterns may further depend on the type of floral reward being evaluated. Whether similar enhancements in nectar reward quantity or quality would generate comparable patterns of response, or even encourage a more liberal acceptance bias, remains unclear. One key aspect of nectar quality - sugar concentration - can be assessed immediately when bees sample nectar (Bitterman, 1976; De Brito Sanchez, 2011; Hemingway & Muth, 2022) and bees can also taste pollen, which may provide some indication of its quality (Mayberry et al., 2024; Muth et al., 2016; Ruedenauer et al., 2016). Future experiments that manipulate the type and magnitude of floral rewards in different ways will be important for testing whether these behavioral patterns generalize across other forms of floral deception.

Aside from the observed asymmetry in decision criteria, we found that learning played an important role in shaping performance over time, with bees adjusting their responses as they became familiar with models and mimics. When mimic costs were increased, bees made proportionally fewer correct decisions and detections overall with experience. In contrast, correct rejections increased with experience regardless of the relative costs or benefits. Buzzing of mimics also diminished with experience when mimic costs were unmodified, indicating bees learned to reduce the energetic cost of false alarms; buzzing of mimics likely did not decline with experience when mimic costs were increased, due to the initially very low incidence of buzzing. Overall, bees showed marginal improvements with experience when models were supplemented with pollen. Pollinator learning is often proposed to mitigate exploitation by floral Batesian mimics (Anderson & Johnson, 2006; Dafni, 1984; Dukas, 1987; Gigord et al., 2001; Jersáková et al., 2006; Schiestl & Johnson, 2013), although the mechanisms underlying this remain poorly understood (de Jager & Ellis, 2014; Gigord et al., 2001, 2002). Naïve pollinators are initially unfamiliar with the phenotypic distribution and payoffs of models and mimics and are therefore expected to initially visit both randomly. With experience they learn to use distinctive cues to target rewarding flowers (Clarke et al., 2013; Rands et al., 2023; Russell & Ashman, 2019; Whitney et al., 2009). Signal detection theory predicts that such learning should lead pollinators to increasingly avoid mimics, even at the cost of some missed rewards, thus maximizing foraging efficiency (Abbott & Sherratt, 2013; Guilford & Dawkins, 1991; Oaten et al., 1975). Our results support this prediction, particularly when mimic costs were high, and align with previous findings that discrimination performance improves with experience (de Jager & Ellis, 2014; Internicola & Harder, 2012; Kunze & Gumbert, 2001; Russell et al., 2020, 2021).

Mimicry occurs across a range of decision-making contexts beyond floral deception, with one of the most well-studied examples involving predators encountering defended prey. In such systems, predators must decide whether to attack or avoid prey that may resemble defended species (Skelhorn et al., 2016). As with pollinators facing deceptive flowers, these decisions entail a trade-off. Being overly cautious may reduce foraging opportunities, whereas being overly liberal may increase risk of exposure to chemically or physically defended prey. Theoretical work has examined how predators evaluate risk in the face of such deceptive signals, exploring how uncertainty and the potential costs of mistakes shape decision rules (Holen & Sherratt, 2021; Kikuchi & Sherratt, 2015; Sherratt, 2011; Speed & Ruxton, 2010). Empirical studies similarly demonstrate that when the costs of attacking defended prey are high, predators tend to adopt more conservative strategies, by avoiding more broadly and sampling less frequently (reviewed in Skelhorn et al., 2016), or relying more heavily on social information (e.g., Hämäläinen et al., 2021). Together, these findings align with the results found in our current study, suggesting that as the costs of errors and the degree of uncertainty increase, decision-makers may favor conservative strategies that minimize potential losses, even at the expense of foregoing greater potential rewards.

In conclusion, bumble bees appear to prioritize avoiding exploitation over maximizing potential gains, revealing an asymmetry in how they evaluate risk when encountering uncertainty with floral mimics. Our findings thus may offer new insight into the evolutionary persistence of floral deception and the ecological dynamics that maintain it. Because pollinators did not substantially increase visits to more profitable models over time, deceptive mimics may persist even in the presence of more rewarding flowers. However, as the costs of visiting deceptive flowers increase, pollinators are expected to adopt even more conservative decision criteria, further reducing visits to potential mimics. If these costs become too high, pollinators may ultimately abandon the entire floral system, decreasing reproductive success for both mimics and models. These dynamics may help explain the stability of automimetic systems like those found in *Begonia*, where deceptive and rewarding flowers coexist with moderate costs (Agren & Schemske, 1991; Schemske et al., 1996; Wyatt & Sazima, 2011). More broadly, a conservative pollinator bias may carry population-level consequences by lowering pollen transfer efficiency and constraining plant reproductive success, thereby shaping the coevolutionary balance between attraction, deception, and learning in plant-pollinator interactions. These dynamics could be tested in both a field and lab setting to shed light on fitness consequences of these decision rules under different cost-benefit regimes. Linking signal detection theory with pollinator behavior in plant–pollinator interactions thus provides a powerful approach for understanding how animals navigate uncertainty and deception in their environments and how these decision-making processes shape ecological and evolutionary outcomes.

## Supporting information

Data and R scripts for analyses

## ACKNOWLEDGEMENTS

We are grateful to Plant Products: Biobest Group for bee colonies, Jenny Burrow for greenhouse care, and Russell lab members for discussion. We acknowledge this work was performed on unceded traditional territory of the Kiikaapoi, Sioux, and Osage.

## DECLARATIONS

## ETHICS APPROVAL

All bumble bee experimentation was carried out in accordance with the legal and ethical standards of the USA.

## DATA ACCESSSIBILITY

The datasets supporting this article are available as electronic supplementary material.

## REFERENCES

Abbott, K. R., & Sherratt, T. N. (2013). Optimal sampling and signal detection: Unifying models of attention and speed–accuracy trade-offs. Behavioral Ecology, 24(3), 605–616. 10.1093/beheco/art001

Agren, J., & Schemske, D. W. (1991). Pollination by Deceit in a Neotropical Monoecious Herb, Begonia involucrata. Biotropica, 23(3), 235. 10.2307/2388200

Anderson, B., & Johnson, S. D. (2006). The effects of floral mimics and models on each others’ fitness. Proceedings of the Royal Society B: Biological Sciences, 273(1589), 969–974. 10.1098/rspb.2005.3401

Barlow, S. E., Wright, G. A., Ma, C., Barberis, M., Farrell, I. W., Marr, E. C., Brankin, A., Pavlik, B. M., & Stevenson, P. C. (2017). Distasteful nectar deters floral robbery. Current Biology, 27(16), 2552–2558.e3. 10.1016/j.cub.2017.07.012

Benadi, G., Kögel, R., Lämsä, J., & Gegear, R. J. (2023). Temporal variation of floral reward can improve the pollination success of a rare flowering plant. Arthropod-Plant Interactions, 17(6), 765–776. 10.1007/s11829-023-10007-8

Bitterman, M. (1976). Incentive contrast in honey bees. Science, 192(4237), 380–382. 10.1126/science.1257773

Blumstein, D. T., & Bouskila, A. (1996). Assessment and decision making in animals: A mechanistic model underlying behavioral flexibility can prevent ambiguity. Oikos, 77, 569–576.

Brooks, M. (2025). glmmTMB: Generalized Linear Mixed Models using Template Model Builder [R].

Cartar, R. V. (2004). Resource tracking by bumble bees: Responses to plant-level differences in quality. Ecology, 85(10), 2764–2771. 10.1890/03-0484

Cartar, RalphV., & Dill, LawrenceM. (1990). Why are bumble bees risk-sensitive foragers? Behavioral Ecology and Sociobiology, 26(2). 10.1007/BF00171581

Chittka, L., Thomson, J. D., & Waser, N. M. (1999). Flower constancy, insect psychology, and plant evolution. Naturwissenschaften, 86(8), 361–377. 10.1007/s001140050636

Clarke, D., Whitney, H., Sutton, G., & Robert, D. (2013). Detection and learning of floral electric fields by bumblebees. Science, 340(6128), 66–69. 10.1126/science.1230883

Cnaani, J., Thomson, J. D., & Papaj, D. R. (2006). Flower choice and learning in foraging bumblebees: Effects of variation in nectar volume and concentration. Ethology, 112(3), 278–285. 10.1111/j.1439-0310.2006.01174.x

Dafni, A. (1984). Mimicry and Deception in Pollination. Annual Review of Ecology and Systematics, 1, 259–278.

De Avila, R. S., Oleques, S. S., Marciniak, B., & Ribeiro, J. R. I. (2017). Effects of model-mimic frequency on insect visitation and plant reproduction in a self-mimicry pollination system. AoB PLANTS, 9(6). 10.1093/aobpla/plx044

De Brito Sanchez, M. G. (2011). Taste Perception in Honey Bees. Chemical Senses, 36(8), 675–692. 10.1093/chemse/bjr040

de Jager, M. L., & Ellis, A. G. (2014). Costs of deception and learned resistance in deceptive interactions. Proceedings of the Royal Society B: Biological Sciences, 281(1779), 20132861. 10.1098/rspb.2013.2861

Developmental Core Team R. (2024). *R: a language and evironment for statistical computing*. R Foundation for Statistical Computing.

Duffy, K. J., & Johnson, S. D. (2017). Effects of distance from models on the fitness of floral mimics. Plant Biology, 19(3), 438–443. 10.1111/plb.12555

Dukas, R. (1987). Foraging behavior of three bee species in a natural mimicry system: Female flowers which mimic male flowers in Ecballium elaterium (Cucurbitaceae). Oecologia, 74(2), 256–263. 10.1007/BF00379368

Ferdy, J., Gouyon, P., Moret, J., & Godelle, B. (1998). Pollinator behavior and deceptive pollination: Learning process and floral evolution. The American Naturalist, 152(5), 696–705.

Ferdy, J.-B., Austerlitz, F., Moret, J., Gouyon, P.-H., & Godelle, B. (1999). Pollinator-induced density dependence in deceptive species. Oikos, 87(3), 549. 10.2307/3546819

Finkbeiner, S. D., Salazar, P. A., Nogales, S., Rush, C. E., Briscoe, A. D., Hill, R. I., Kronforst, M. R., Willmott, K. R., & Mullen, S. P. (2018). Frequency dependence shapes the adaptive landscape of imperfect Batesian mimicry. Proceedings of the Royal Society B: Biological Sciences, 285(1876), 20172786. 10.1098/rspb.2017.2786

Fox, J., Weisberg, S., Adler, D., Bates, D., Baud-Bovy, G., &, et al. (2023). *Package “car”* (Version 3.1-2) [R Core Language].

Gaskett, A. C. (2011). Orchid pollination by sexual deception: Pollinator perspectives. Biological Reviews, 86(1), 33–75. 10.1111/j.1469-185X.2010.00134.x

Gigord, L. D. B., Macnair, M. R., & Smithson, A. (2001). Negative frequency-dependent selection maintains a dramatic flower color polymorphism in the rewardless orchid *Dactylorhiza sambucina* (L.) Soò. Proceedings of the National Academy of Sciences, 98(11), 6253–6255. 10.1073/pnas.111162598

Gigord, L. D. B., Macnair, M. R., Stritesky, M., & Smithson, A. (2002). The potential for floral mimicry in rewardless orchids: An experimental study. Proceedings of the Royal Society of London. Series B: Biological Sciences, 269(1498), 1389–1395. 10.1098/rspb.2002.2018

Green, D., & Swets, J. (1966). Signal detection theory and psychophysics (Vol. 1). Wiley.

Guilford, T., & Dawkins, M. S. (1991). Receiver psychology and the evolution of animal signals. Animal Behaviour, 42(1), 1–14.

Gumbert, A., & Kunze, J. (2001). Colour similarity to rewarding model plants affects pollination in a food deceptive orchid, Orchis boryi. Biological Journal of the Linnean Society, 72(3), 419–433. 10.1111/j.1095-8312.2001.tb01328.x

Hämäläinen, L., Hoppitt, W., Rowland, H. M., Mappes, J., Fulford, A. J., Sosa, S., & Thorogood, R. (2021). Social transmission in the wild can reduce predation pressure on novel prey signals. Nature Communications, 12(1), 3978. 10.1038/s41467-021-24154-0

Harder, L. D., & Real, L. A. (1987). Why are bumble bees risk averse? Ecology, 68(4), 1104–1108. 10.2307/1938384

Hartig, F. (2024). DHARMa: Residual Diagnostics for Hierarchical (Multi-level/Mixed) Regression Models (Version 0.4.7) [R].

Heinrich, B. (1976). The foraging specializations of individual bumblebees. Ecological Monographs, 46, 105–128.

Heinrich, B. (1979). *Bumblebee Economics* (6th ed.). Harvard University Press.

Hemingway, C. T., DeVore, J. E., & Muth, F. (2024). Economic foraging in a floral marketplace: Asymmetrically dominated decoy effects in bumblebees. Proceedings of the Royal Society B: Biological Sciences, 291(2031), 20240843. 10.1098/rspb.2024.0843

Hemingway, C. T., Leonard, A. S., MacNeill, F. T., Pimplikar, S., & Muth, F. (2024). Pollinator cognition and the function of complex rewards. Trends in Ecology & Evolution, 39(11), 1047–1058. 10.1016/j.tree.2024.06.008

Hemingway, C. T., & Muth, F. (2022). Label-based expectations affect incentive contrast effects in bumblebees. Biology Letters, 18, 20210549.

Heyneman, A. J. (1983). Optimal sugar concentrations of floral nectars—Dependence on sugar intake efficiency and foraging costs. Oecologia, 60(2), 198–213. 10.1007/BF00379522

Holen, Ø. H., & Sherratt, T. N. (2021). Coping with Danger and Deception: Lessons from Signal Detection Theory. The American Naturalist, 197(2), 147–163. 10.1086/712246

Internicola, A. I., & Harder, L. D. (2012). Bumble-bee learning selects for both early and long flowering in food-deceptive plants. Proceedings of the Royal Society B: Biological Sciences, 279(1733), 1538–1543. 10.1098/rspb.2011.1849

Jersáková, J., Johnson, S. D., & Kindlmann, P. (2006). Mechanisms and evolution of deceptive pollination in orchids. Biological Reviews, 81(2), 219–235. 10.1017/S1464793105006986

Johnson, S., & Schiestl, F. P. (2016). Floral mimicry. Oxford University Press.

Jones, P. L., & Agrawal, A. A. (2016). Consequences of toxic secondary compounds in nectar for mutualist bees and antagonist butterflies. Ecology, 97(10), 2570–2579. 10.1002/ecy.1483

Kacelnik, A., & Bateson, M. (1996). Risky Theories—The Effects of Variance on Foraging Decisions. American Zoologist, 36(4), 402–434. 10.1093/icb/36.4.402

Kikuchi, D. W., & Sherratt, T. N. (2015). Costs of Learning and the Evolution of Mimetic Signals. The American Naturalist, 186(3), 321–332. 10.1086/682371

Kunze, J., & Gumbert, A. (2001). The combined effect of color and odor on flower choice behavior of bumble bees in flower mimicry systems. Behavioral Ecology, 12(4), 447–456. 10.1093/beheco/12.4.447

Length, R. (2025). emmeans: Estimated Marginal Means, aka Least-Squares Means [Computer software].

Leonard, A. S., Dornhaus, A., & Papaj, D. R. (2011). Flowers help bees cope with uncertainty: Signal detection and the function of floral complexity. Journal of Experimental Biology, 214(1), 113–121. 10.1242/jeb.047407

Lichtenberg, E. M., Heiling, J. M., Bronstein, J. L., & Barker, J. L. (2020). Noisy communities and signal detection: Why do foragers visit rewardless flowers? Philosophical Transactions of the Royal Society B: Biological Sciences, 375(1802), 20190486. 10.1098/rstb.2019.0486

Lindström, L., Alatalo, R. V., & Mappes, J. (1997). Imperfect Batesian mimicry—The effects of the frequency and the distastefulness of the model. Proceedings of the Royal Society of London. Series B: Biological Sciences, 264(1379), 149–153. 10.1098/rspb.1997.0022

Lynn, S. K., & Barrett, L. F. (2014). “Utilizing” signal detection theory. Psychological Science, 25(9), 1663–1673. 10.1177/0956797614541991

Mayberry, M. M., Naumer, K. C., Novinger, A. N., McCart, D. M., Wilkins, R. V., Muse, H., Ashman, T.-L., & Russell, A. L. (2024). Learning to handle flowers increases pollen collection for bees but does not affect pollination success for plants. Behavioral Ecology, 35(6), arae083. 10.1093/beheco/arae083

McNamara, J. M., & Houston, A. I. (1992). Risk-sensitive foraging: A review of the theory. Bulletin of Mathematical Biology, 54, 355–378.

Muth, F., Francis, J. S., & Leonard, A. S. (2016). Bees use the taste of pollen to determine which flowers to visit. Biology Letters, 12(7), 20160356. 10.1098/rsbl.2016.0356

Nachev, V., Stich, K. P., Winter, C., Bond, A., Kamil, A., & Winter, Y. (2017). Cognition-mediated evolution of low-quality floral nectars. Science, 355(6320), 75–78. 10.1126/science.aah4219

Nakazawa, T., Matsumoto, T. K., & Katsuhara, K. R. (2024). When is lethal deceptive pollination maintained? A population dynamics approach. Annals of Botany, 134(4), 665–682. 10.1093/aob/mcae108

Nicholls, E., & Hempel De Ibarra, N. (2017). Assessment of pollen rewards by foraging bees. Functional Ecology, 31(1), 76–87. 10.1111/1365-2435.12778

Oaten, A., Pearce, C. E. M., & Smyth, M. E. B. (1975). Batesian mimicry and signal detection theory. Bulletin of Mathematical Biology, 37, 367–387.

Pannell, J. R., & Farmer, E. E. (2016). Mimicry in plants. Current Biology, 26(17), R784–R785. 10.1016/j.cub.2016.04.005

Pemberton, R. W., & Wheeler, G. S. (2006). Orchid bees don’t need orchids: Evidence from the naturalization of an orchid bee in Florida. Ecology, 87(8), 1995–2001. 10.1890/0012-9658(2006)87%255B1995:OBDNOE%255D2.0.CO;2

Pfennig, D. W., Harcombe, W. R., & Pfennig, K. S. (2001). Frequency-dependent Batesian mimicry. Nature, 410(6826), 323–323. 10.1038/35066628

Pyke, G. H., & Ren, Z. (2023). Floral nectar production: What cost to a plant? Biological Reviews, 98(6), 2078–2090. 10.1111/brv.12997

Raine, N. E., & Chittka, L. (2008). The correlation of learning speed and natural foraging success in bumblebees. Proceedings of the Royal Society B: Biological Sciences, 275, 803–808.

Rands, S. A., Whitney, H. M., & Hempel De Ibarra, N. (2023). Multimodal floral recognition by bumblebees. Current Opinion in Insect Science, 59, 101086. 10.1016/j.cois.2023.101086

Real, L. A. (1992). Information Processing and the Evolutionary Ecology of Cognitive Architecture. The American Naturalist, 140, S108–S145. 10.1086/285399

Renner, S. (2006). Rewardless flowers in the angiosperms and the role of insect cognition in their evolution. In Plant-pollinator interactions: From specialization to generalization (pp. 123–144).

Richman, S. K., Irwin, R. E., & Bronstein, J. L. (2017). Foraging strategy predicts foraging economy in a facultative secondary nectar robber. Oikos, 126(9), 1250–1257. 10.1111/oik.04229

Robertson, A. W., Mountjoy, C., Faulkner, B. E., Roberts, M. V., & Macnair, M. R. (1999). Bumble bee selection of *Mimulus guttatus* flowers: The effects of pollen quality and reward depletion. Ecology, 80(8), 2594–2606. 10.1890/0012-9658(1999)080%255B2594:BBSOMG%255D2.0.CO;2

Romero, G. Q., Antiqueira, P. A. P., & Koricheva, J. (2011). A meta-analysis of predation risk effects on pollinator behaviour. PLoS ONE, 6(6), e20689. 10.1371/journal.pone.0020689

Rossi, N., Vallejo-Marín, M., & Nicholls, E. (2025). First direct quantification of floral handling costs in bees. bioRxiv. 10.1101/2025.09.15.675838

Ruedenauer, F. A., Spaethe, J., & Leonhardt, S. D. (2016). Hungry for quality—Individual bumblebees forage flexibly to collect high-quality pollen. Behavioral Ecology and Sociobiology, 70(8), 1209–1217. 10.1007/s00265-016-2129-8

Russell, A. L., & Ashman, T.-L. (2019). Associative learning of flowers by generalist bumble bees can be mediated by microbes on the petals. Behavioral Ecology, 30(3), 746–755. 10.1093/beheco/arz011

Russell, A. L., Buchmann, S. L., & Papaj, D. R. (2017). How a generalist bee achieves high efficiency of pollen collection on diverse floral resources. Behavioral Ecology, 28(4), 991–1003. 10.1093/beheco/arx058

Russell, A. L., Kikuchi, D. W., Giebink, N. W., & Papaj, D. R. (2020). Sensory bias and signal detection trade-offs maintain intersexual floral mimicry. Philosophical Transactions of the Royal Society B: Biological Sciences, 375(1802), 20190469. 10.1098/rstb.2019.0469

Russell, A. L., Sanders, S. R., Wilson, L. A., & Papaj, D. R. (2021). The size of it: Scant evidence that flower size variation affects deception in intersexual floral mimicry. Frontiers in Ecology and Evolution, 9, 724712. 10.3389/fevo.2021.724712

Schemske, D. W., Ågren, J., & Corff, J. L. (1996). Deceit Pollination in the Monoecious, Neotropical Herb Begonia oaxacana (Begoniaceae). In D. G. Lloyd & S. C. H. Barrett (Eds.), Floral Biology (pp. 292–318). Springer US. 10.1007/978-1-4613-1165-2_11

Schiestl, F. P., & Johnson, S. D. (2013). Pollinator-mediated evolution of floral signals. Trends in Ecology & Evolution, 28(5), 307–315. 10.1016/j.tree.2013.01.019

Shafir, S., Reich, T., Tsur, E., Erev, I., & Lotem, A. (2008). Perceptual accuracy and conflicting effects of certainty on risk-taking behaviour. Nature, 453(7197), 917–920. 10.1038/nature06841

Sherratt, T. N. (2011). The optimal sampling strategy for unfamiliar prey. Evolution, 65(7), 2014–2025. 10.1111/j.1558-5646.2011.01274.x

Sherratt, T. N., & Peet-Paré, C. A. (2017). The perfection of mimicry: An information approach. Philosophical Transactions of the Royal Society B: Biological Sciences, 372(1724), 20160340. 10.1098/rstb.2016.0340

Signorell, A., Aho, K., Alfons, A., Anderegg, N., Aragon, T., Arachchige, C., Arppe, A., &, et al. (2025). DescTools: Tools for Descriptive Statistics [R].

Skelhorn, J., Halpin, C. G., & Rowe, C. (2016). Learning about aposematic prey. Behavioral Ecology, 27(4), 955–964. 10.1093/beheco/arw009

Southwick, E. E. (1984). Photosynthate allocation to floral nectar: A neglected energy investment. Ecology, 65(6), 1775–1779. 10.2307/1937773

Speed, M. P., & Ruxton, G. D. (2010). Imperfect Batesian Mimicry and the Conspicuousness Costs of Mimetic Resemblance. The American Naturalist, 176(1), E1–E14. 10.1086/652990

Stephens, D. W. (2008). Decision ecology: Foraging and the ecology of animal decision making. Cognitive, Affective, & Behavioral Neuroscience, 8, 475–484.

Stevenson, P. C., Nicolson, S. W., & Wright, G. A. (2017). Plant secondary metabolites in nectar: Impacts on pollinators and ecological functions. Functional Ecology, 31(1), 65–75. 10.1111/1365-2435.12761

Vázquez, V., & Barradas, I. (2017). Deceptive pollination and insects’ learning: A delicate balance. Journal of Biological Dynamics, 11(1), 299–322. 10.1080/17513758.2017.1337246

Venjakob, C., Ruedenauer, F. A., Klein, A. M., & Leonhardt, S. D. (2022). Variation in nectar quality across 34 grassland plant species. Plant Biology, 24(1), 134–144. 10.1111/plb.13343

Whitney, H. M., Chittka, L., Bruce, T. J. A., & Glover, B. J. (2009). Conical epidermal cells allow bees to grip flowers and increase foraging efficiency. Current Biology, 19(11), 948–953. 10.1016/j.cub.2009.04.051

Wiley, R. H. (2006). Signal detection and animal communication. In Advances in the Study of Behavior (Vol. 36, pp. 217–247). Elsevier. 10.1016/S0065-3454(06)36005-6

Wolf, L. L., Hainsworth, F. R., & Gill, F. B. (1975). Foraging efficiencies and time budgets in nectar-feeding birds. Ecology, 56(1), 117–128. 10.2307/1935304

Wyatt, G. E., & Sazima, M. (2011). Pollination and reproductive biology of thirteen species of *Begonia* in the Serra do Mar State Park, São Paulo, Brazil. Journal of Pollination Ecology, 6. 10.26786/1920-7603(2011)16

Zurbuchen, A., Cheesman, S., Klaiber, J., Müller, A., Hein, S., & Dorn, S. (2010). Long foraging distances impose high costs on offspring production in solitary bees. Journal of Animal Ecology, 79(3), 674–681. 10.1111/j.1365-2656.2010.01675.x

