## Supplementary material for "Bees attend primarily to costs, not benefits, to avoid exploitation by floral mimics": Data and R scripts for analyses: Stats_60_SDT-costs-benefits_5-29-2024.R

```
# Changelog ####
# 4-11-2023 File created
# 5-4-2023 ANN truncated everything to 60 visits
# 5-5-2023 ALR modified plots and cleaned up code
# 5-5-2023 ALR added modified/unmodified code 
# 5-9-2023 ANN fixed comments for questions 8-10 (were previously unaltered from
#          being copied from questions 2-4), switched the figure names for 
#          questions 5 and 6 (switched from 2c,2b to 2b,2c, respectively) to 
#          match the order of the questions.
# 5-10-2023 ALR added headers (yay!!) and fixed convergence error in Q4
#           ANN updated some of the commented stats results (Q4 and Q5)          


#Name of data file:
FILENAME1 <- "SDT_60_CorrectDecisions_5-4_23_AN.csv"
FILENAME2 <- "SDT_60_CorrectDetections_5-4-23_AN.csv"
FILENAME3 <- "SDT_60_CorrectRejections_5-4-23_AN.csv"
FILENAME4 <- "SDT_60_LandsOnMale_5-4-23_AN.csv"
FILENAME5 <- "SDT_60_BuzzGivenLand_Male_5-4-23_AN.csv"
FILENAME6 <- "SDT_60_BuzzGivenLand_Female_5-4-23_AN.csv"
FILENAME7 <- "SDT_60_UnmodVsModControl_CorrectDecisions_5-4-23_AN.csv"
FILENAME8 <- "SDT_60_UnmodVsModControl_CorrectDetections_5-4-23_AN.csv"
FILENAME9 <- "SDT_60_UnmodVsModControl_CorrectRejections_5-4-23_AN.csv"
FILENAME10 <- "SDT_60_UnmodVsModControl_LandsOnMale_5-4-23_AN.csv"
#FILENAME <- ".csv"

#Sets working directory automatically to where R script is saved
setwd(dirname(rstudioapi::getActiveDocumentContext()$path))

# Set the directory your plots will be in:
PLOTDIR <- dirname(rstudioapi::getActiveDocumentContext()$path)

#Load data:
setwd(dirname(rstudioapi::getActiveDocumentContext()$path)); dataset1 <- read.table(FILENAME1, sep=",", header=TRUE)
setwd(dirname(rstudioapi::getActiveDocumentContext()$path)); dataset2 <- read.table(FILENAME2, sep=",", header=TRUE)
setwd(dirname(rstudioapi::getActiveDocumentContext()$path)); dataset3 <- read.table(FILENAME3, sep=",", header=TRUE)
setwd(dirname(rstudioapi::getActiveDocumentContext()$path)); dataset4 <- read.table(FILENAME4, sep=",", header=TRUE)
setwd(dirname(rstudioapi::getActiveDocumentContext()$path)); dataset5 <- read.table(FILENAME5, sep=",", header=TRUE)
setwd(dirname(rstudioapi::getActiveDocumentContext()$path)); dataset6 <- read.table(FILENAME6, sep=",", header=TRUE)
setwd(dirname(rstudioapi::getActiveDocumentContext()$path)); dataset7 <- read.table(FILENAME7, sep=",", header=TRUE)
setwd(dirname(rstudioapi::getActiveDocumentContext()$path)); dataset8 <- read.table(FILENAME8, sep=",", header=TRUE)
setwd(dirname(rstudioapi::getActiveDocumentContext()$path)); dataset9 <- read.table(FILENAME9, sep=",", header=TRUE)
setwd(dirname(rstudioapi::getActiveDocumentContext()$path)); dataset10 <- read.table(FILENAME10, sep=",", header=TRUE)

#libraries
library("mgcv")
library("lmerTest")
library("mclust")
library("DescTools")
library("lme4")
library("lsmeans")
library("multcomp")
library("DHARMa") #Checking model assumptions
library("sjPlot") #Checking model assumptions
library("car")
library("glmmTMB")
library("effects") #Making plots
library("ggplot2") #Making plots
library("RVAideMemoire")
library("tidyverse")
library("hrbrthemes")
library("viridis") #Making plots


#_______QUESTION SCRIPTS BELOW_______#

#_______________________________________________________________________________________________________________________________#
# QUESTION 1 ####
# Do bees learn differently when there's different costs and benefits of mimics and models?
# Did bee learn to make CORRECT DECISIONS? Analyze all visits to male and female together.

# Run the model
# Nested random effects; NumVisits within Bee within Colony
All.Correct <- glmmTMB(Correct ~ Treatment * VisitNum + (1 | ColonyID / BeeID / VisitNum), family = binomial, data = dataset1)

# Examine the model
summary(All.Correct)

# Check Type II sums of squares, for Wald Tests
Anova(All.Correct)

# Response: Correct
#                        Chisq Df Pr(>Chisq)    
# Treatment          19.7064  3  0.0001953 ***
# VisitNum            6.3767  1  0.0115625 *  
# Treatment:VisitNum 10.0553  3  0.0181015 *

# Post-hoc test
emmeans(All.Correct, list(pairwise ~ Treatment*VisitNum), adjust = "tukey")

# Pairwise differences among Treatments
#                           estimate     SE   df t.ratio p.value
# Inc_Benefit - Inc_Both       0.432 0.1038 3913   4.161  0.0002
# Inc_Benefit - Inc_Cost       0.317 0.1144 3913   2.775  0.0284
# Inc_Benefit - Mod_Control    0.103 0.0991 3913   1.038  0.7274
# Inc_Both - Inc_Cost         -0.115 0.1181 3913  -0.971  0.7658
# Inc_Both - Mod_Control      -0.329 0.1034 3913  -3.183  0.0080
# Inc_Cost - Mod_Control      -0.214 0.1140 3913  -1.882  0.2360

# Model assumptions via DHARMa
simulationOutput1 <- simulateResiduals(fittedModel = All.Correct, n = 1000)
plot(simulationOutput1) #Plot of QQ plot looks good
testUniformity(simulationOutput = simulationOutput1) # Uniformity good
testDispersion(simulationOutput1) # Dispersion good
testZeroInflation(simulationOutput1) #Zero inflation good

# Model assumptions via sjPlot
# Normality of residuals and outliers
plot_model(All.Correct, type = "diag") #Looks normal

## Plot the model ####
# Standardized visit number
effect1 <- Effect(focal.predictors=c("Treatment","VisitNum"),All.Correct)
effect1 <- as.data.frame(effect1)

#Rename the figure legends
effect1$Treatment = plyr::mapvalues(effect1$Treatment, from = c("Inc_Benefit", "Inc_Cost", "Inc_Both", "Mod_Control"), 
                                     to = c("Benefit", "Cost","Both","Control"))

# Plot model with 95% CI #
Figure1a <- ggplot(effect1, aes(x=VisitNum, y=fit, color=Treatment, fill=Treatment)) + 
  geom_line(lwd=1) +
  geom_ribbon(aes(ymin=lower, ymax=upper), alpha=0.2, colour=NA) + 
  scale_linetype_manual(values=c("solid","solid","solid")) + 
  ylab("Proportion correct decisions") + xlab("Visit number") + theme_classic() + 
  theme(text=element_text(size=14)) +
  scale_color_viridis_d(name = "Treatment", end = 0.7, option = "inferno") + 
  scale_fill_viridis_d(name = "Treatment", end =0.7, option = 'inferno') +
  theme(legend.position  =c(0.5, 0.9), legend.direction = "horizontal", 
        legend.background = element_rect(fill="transparent", colour = NA),
        strip.background = element_blank(),
        legend.title = element_text(size = 10), legend.text = element_text(size = 10),
        panel.spacing.x = unit(1, "lines"), 
        plot.margin = margin(c(0.2,0.51, 0.2, 0.2), unit = "cm")) + 
        guides(colour=guide_legend(nrow=2, byrow=TRUE)) +
  xlim(0,60) + ylim(0,1)
plot(Figure1a) #To see the figure in R

# Set date
DATE <- "5-5-2023"

# Figure dimensions and name
tiff(filename=paste("Fig1a_CorrectDecisions_60vis_", DATE, ".tif", sep=""), units="mm", width=85, height=85, res=1200)

# Export the figure
plot(Figure1a)

# Close the plotting device
setwd(PLOTDIR); 
dev.off()

#_______________________________________________________________________________________________________________________________#


#_______________________________________________________________________________________________________________________________#
# QUESTION 2 ####
# Do bees learn differently when there's different costs and benefits of mimics and models?
# Did bee learn to make CORRECT DETECTIONS? Analyze only visits to male.

# Run the model
# Nested random effects; VisitNum within Bee within Colony
Male.Correct <- glmmTMB(Correct ~ Treatment * VisitNum + (1 | ColonyID / BeeID / VisitNum), family = binomial, data = dataset2)

# Examine the model
summary(Male.Correct)

# Check Type II sums of squares, for Wald Tests
Anova(Male.Correct)

# Response: Correct
#                      Chisq Df Pr(>Chisq)    
# Treatment          45.7886  3   6.29e-10 ***
# VisitNum           96.9680  1  < 2.2e-16 ***
# Treatment:VisitNum  8.5689  3    0.03561 

# Post-hoc test
emmeans(Male.Correct, list(pairwise ~ Treatment), adjust = "tukey")

#$`pairwise differences of Treatment`
#1                         estimate    SE  df z.ratio p.value
# Inc_Benefit - Inc_Both      1.5939 0.332 Inf   4.805  <.0001
# Inc_Benefit - Inc_Cost      1.4983 0.343 Inf   4.372  0.0001
# Inc_Benefit - Mod_Control  -0.1659 0.343 Inf  -0.484  0.9626
# Inc_Both - Inc_Cost        -0.0956 0.333 Inf  -0.288  0.9917
# Inc_Both - Mod_Control     -1.7598 0.338 Inf  -5.205  <.0001
# Inc_Cost - Mod_Control     -1.6641 0.349 Inf  -4.768  <.0001

# Model assumptions via DHARMa
simulationOutput1 <- simulateResiduals(fittedModel = Male.Correct, n = 1000)
plot(simulationOutput1) # QQ plot has a significant deviation 0.01
testUniformity(simulationOutput = simulationOutput1) # Uniformity good
testDispersion(simulationOutput1) # Dispersion good
testZeroInflation(simulationOutput1) #Zero inflation good

# Model assumptions via sjPlot
# Normality of residuals and outliers
plot_model(Male.Correct, type = "diag") #

## Plot the model ####
# Standardized visit number
effect2 <- Effect(focal.predictors=c("Treatment","VisitNum"),Male.Correct)
effect2 <- as.data.frame(effect2)

#Rename the figure legends
effect2$Treatment = plyr::mapvalues(effect2$Treatment, from = c("Inc_Benefit", "Inc_Cost", "Inc_Both", "Mod_Control"), 
                                    to = c("Benefit", "Cost","Both","Control"))

# Plot model with 95% CI
Figure1b <- ggplot(effect2, aes(x=VisitNum, y=fit, color=Treatment, fill=Treatment)) + 
  geom_line(lwd=1) +
  geom_ribbon(aes(ymin=lower, ymax=upper), alpha=0.2, colour=NA) + 
  scale_linetype_manual(values=c("solid","solid","solid")) + 
  ylab("Proportion correct detections") + xlab("Visit number") + theme_classic() + 
  theme(text=element_text(size=14)) +
  scale_color_viridis_d(name = "Treatment", end = 0.7, option = "inferno") + 
  scale_fill_viridis_d(name = "Treatment", end =0.7, option = 'inferno') +
  theme(legend.position  =c(0.5, 0.1), legend.direction = "horizontal", 
        legend.background = element_rect(fill="transparent", colour = NA),
        strip.background = element_blank(),
        legend.title = element_text(size = 10), legend.text = element_text(size = 10),
        panel.spacing.x = unit(1, "lines"), 
        plot.margin = margin(c(0.2,0.51, 0.2, 0.2), unit = "cm")) + 
        guides(colour=guide_legend(nrow=2, byrow=TRUE)) +
  xlim(0,60) + ylim(0,1)
plot(Figure1b) #To see the figure in R

# Set date
DATE <- "5-5-2023"

# Figure dimensions and name
tiff(filename=paste("Fig1b_CorrectDetections_60vis_", DATE, ".tif", sep=""), units="mm", width=85, height=85, res=1200)

# Export the figure
plot(Figure1b)

# Close the plotting device
setwd(PLOTDIR); 
dev.off()

#_______________________________________________________________________________________________________________________________#


#_______________________________________________________________________________________________________________________________#
# QUESTION 3 ####
# Do bees learn differently when there's different costs and benefits of mimics and models?
# Did bee learn to make CORRECT REJECTIONS? Analyze only visits to female. 
# Run the model


# Nested random effects; NumVisits within Bee
# Model does not converge with ColonyID
Female.Correct <- glmmTMB(Correct ~ Treatment * VisitNum + (1 | BeeID / VisitNum), family = binomial, data = dataset3)

# Examine the model
summary(Female.Correct)

# Check Type II sums of squares, for Wald Tests
Anova(Female.Correct)

#Response: Correct
#                       Chisq Df Pr(>Chisq)    
# Treatment            6.9385  3    0.07389 .  
# VisitNum           130.0709  1    < 2e-16 ***
# Treatment:VisitNum   0.6715  3    0.87989 

# Post-hoc test
emmeans(Female.Correct, list(pairwise ~ Treatment), adjust = "tukey")

#$`pairwise differences of Treatment`
#1                         estimate    SE  df z.ratio p.value
# Inc_Benefit - Inc_Both      -0.158 0.225 2021  -0.702  0.8964
# Inc_Benefit - Inc_Cost      -0.341 0.240 2021  -1.417  0.4888
# Inc_Benefit - Mod_Control    0.242 0.217 2021   1.114  0.6808
# Inc_Both - Inc_Cost         -0.183 0.245 2021  -0.745  0.8787
# Inc_Both - Mod_Control       0.399 0.225 2021   1.773  0.2869
# Inc_Cost - Mod_Control       0.582 0.241 2021   2.415  0.0745

# Model assumptions via DHARMa
simulationOutput1 <- simulateResiduals(fittedModel = Female.Correct, n = 1000)
plot(simulationOutput1) # QQ plot is good
testUniformity(simulationOutput = simulationOutput1) # Uniformity good
testDispersion(simulationOutput1) # Dispersion good
testZeroInflation(simulationOutput1) #Zero inflation good

# Model assumptions via sjPlot
# Normality of residuals and outliers
plot_model(Female.Correct, type = "diag") #

## Plot the model ####
# Standardized visit number
effect3 <- Effect(focal.predictors=c("Treatment","VisitNum"),Female.Correct)
effect3 <- as.data.frame(effect3)

#Rename the figure legends
effect3$Treatment = plyr::mapvalues(effect3$Treatment, from = c("Inc_Benefit", "Inc_Cost", "Inc_Both", "Mod_Control"), 
                                    to = c("Benefit", "Cost","Both","Control"))

# Plot model with 95% CI
Figure1c <- ggplot(effect3, aes(x=VisitNum, y=fit, color=Treatment, fill=Treatment)) + 
  geom_line(lwd=1) +
  geom_ribbon(aes(ymin=lower, ymax=upper), alpha=0.2, colour=NA) + 
  scale_linetype_manual(values=c("solid","solid","solid")) + 
  ylab("Proportion correct rejections") + xlab("Visit number") + theme_classic() + 
  theme(text=element_text(size=14)) +
  scale_color_viridis_d(name = "Treatment", end = 0.7, option = "inferno") + 
  scale_fill_viridis_d(name = "Treatment", end =0.7, option = 'inferno') +
  theme(legend.position  =c(0.5, 0.92), legend.direction = "horizontal", 
        legend.background = element_rect(fill="transparent", colour = NA),
        strip.background = element_blank(),
        legend.title = element_text(size = 10), legend.text = element_text(size = 10),
        panel.spacing.x = unit(1, "lines"), 
        plot.margin = margin(c(0.2,0.51, 0.2, 0.2), unit = "cm")) + 
        guides(colour=guide_legend(nrow=2, byrow=TRUE)) +
  xlim(0,60) + ylim(0,1)
plot(Figure1c) #To see the figure in R

# Set date
DATE <- "5-5-2023"

# Figure dimensions and name
tiff(filename=paste("Fig1c_CorrectRejections_60vis_", DATE, ".tif", sep=""), units="mm", width=85, height=85, res=1200)

# Export the figure
plot(Figure1c)

# Close the plotting device
setwd(PLOTDIR); 
dev.off()
#_______________________________________________________________________________________________________________________________#


#_______________________________________________________________________________________________________________________________#
# QUESTION 4 ####
# Analyze all LANDS, proportion that are on males vs females.

# Run the model
# Nested random effects; NumVisits within Bee within Colony
All.Male.Land <- glmmTMB(LandsOnMale ~ Treatment * VisitNum + (1 | ColonyID / BeeID / VisitNum), family = binomial, data = dataset4)

# Examine the model
summary(All.Male.Land)

# Check Type II sums of squares, for Wald Tests
Anova(All.Male.Land)

#Response: LandsOnMale
#                    Chisq Df Pr(>Chisq)  
#Treatment          4.9349  3    0.17663  
# VisitNum           2.9626  1    0.08521 .
# Treatment:VisitNum 8.4066  3    0.03832 *

# Post-hoc test
emmeans(All.Male.Land, list(pairwise ~ Treatment*VisitNum), adjust = "tukey")

#$`pairwise differences of Treatment`
#1                         estimate    SE  df z.ratio p.value
# Inc_Benefit - Inc_Both      0.2483 0.114 2599   2.188  0.1269
# Inc_Benefit - Inc_Cost      0.1707 0.123 2599   1.385  0.5089
# Inc_Benefit - Mod_Control   0.1003 0.101 2599   0.993  0.7535
# Inc_Both - Inc_Cost        -0.0775 0.133 2599  -0.585  0.9368
# Inc_Both - Mod_Control     -0.1480 0.112 2599  -1.318  0.5515
# Inc_Cost - Mod_Control     -0.0704 0.122 2599  -0.577  0.9391

# Model assumptions via DHARMa
simulationOutput1 <- simulateResiduals(fittedModel = All.Male.Land, n = 1000)
plot(simulationOutput1) # QQ plot is good
testUniformity(simulationOutput = simulationOutput1) # Uniformity good
testDispersion(simulationOutput1) # Dispersion good
testZeroInflation(simulationOutput1) #Zero inflation good

# Model assumptions via sjPlot
# Normality of residuals and outliers
plot_model(All.Male.Land, type = "diag") #


## Plot the model ####
# Standardized visit number
effect5 <- Effect(focal.predictors=c("Treatment","VisitNum"),All.Male.Land)
effect5 <- as.data.frame(effect5)

#Rename the figure legends
effect5$Treatment = plyr::mapvalues(effect5$Treatment, from = c("Inc_Benefit", "Inc_Cost", "Inc_Both", "Mod_Control"), 
                                    to = c("Benefit", "Cost","Both","Control"))

# Plot model with 95% CI
Figure2a <- ggplot(effect5, aes(x=VisitNum, y=fit, color=Treatment, fill=Treatment)) + 
  geom_line(lwd=1) +
  geom_ribbon(aes(ymin=lower, ymax=upper), alpha=0.2, colour=NA) + 
  scale_linetype_manual(values=c("solid","solid","solid")) + 
  ylab("Proportion lands on models") + xlab("Visit number") + theme_classic() + 
  theme(text=element_text(size=14)) +
  scale_color_viridis_d(name = "Treatment", end = 0.7, option = "inferno") + 
  scale_fill_viridis_d(name = "Treatment", end =0.7, option = 'inferno') +
  theme(legend.position  =c(0.5, 0.9), legend.direction = "horizontal", 
        legend.background = element_rect(fill="transparent", colour = NA),
        strip.background = element_blank(),
        legend.title = element_text(size = 10), legend.text = element_text(size = 10),
        panel.spacing.x = unit(1, "lines"), 
        plot.margin = margin(c(0.2,0.51, 0.2, 0.2), unit = "cm")) + 
        guides(colour=guide_legend(nrow=2, byrow=TRUE)) +
  xlim(0,60) + ylim(0,1)
plot(Figure2a) #To see the figure in R

# Set date
DATE <- "5-5-2023"

# Figure dimensions and name
tiff(filename=paste("Fig2a_ProportionLandsOnMale_60vis_", DATE, ".tif", sep=""), units="mm", width=85, height=85, res=1200)

# Export the figure
plot(Figure2a)

# Close the plotting device
setwd(PLOTDIR); 
dev.off()
#_______________________________________________________________________________________________________________________________#


#_______________________________________________________________________________________________________________________________#
# QUESTION 5 ####
# Do bees BUZZ on males more frequently with experience and does this depend on different adjustments of costs and benefits?
# Analyze only MALE.

# Run the model
# Nested random effects; NumVisits within Bee within colony
# QQ plot is not normal when ColonyID included
Buzz.Given.Land.Male <- glmmTMB(BuzzGivenLand ~ Treatment * VisitNum + (1 | BeeID / VisitNum), family = binomial, data = dataset5)

# Examine the model
summary(Buzz.Given.Land.Male)

# Check Type II sums of squares, for Wald Tests
Anova(Buzz.Given.Land.Male)

#Response: BuzzGivenLand
#                     Chisq Df Pr(>Chisq)    
#Treatment          66.4856  3  2.413e-14 ***
#VisitNum            1.5568  1     0.2121    
#Treatment:VisitNum  3.5664  3     0.3122   

# Post-hoc test
emmeans(Buzz.Given.Land.Male, list(pairwise ~ Treatment), adjust = "tukey")

# $`pairwise differences of Treatment`
# 1                         estimate    SE   df t.ratio p.value
# Inc_Benefit - Inc_Both       3.382 0.640 1420   5.286  <.0001
# Inc_Benefit - Inc_Cost       3.499 0.661 1420   5.295  <.0001
# Inc_Benefit - Mod_Control   -1.480 0.705 1420  -2.099  0.1539
# Inc_Both - Inc_Cost          0.117 0.632 1420   0.186  0.9977
# Inc_Both - Mod_Control      -4.862 0.729 1420  -6.668  <.0001
# Inc_Cost - Mod_Control      -4.980 0.752 1420  -6.620  <.0001

# Model assumptions via DHARMa
simulationOutput1 <- simulateResiduals(fittedModel = Buzz.Given.Land.Male, n = 1000)
plot(simulationOutput1) # QQ plot is good
testUniformity(simulationOutput = simulationOutput1) # Uniformity good 
testDispersion(simulationOutput1) # Dispersion good
testZeroInflation(simulationOutput1) # Problems ratioObsSim = 0.76564, p-value = 0.016

# Model assumptions via sjPlot
# Normality of residuals and outliers
plot_model(Buzz.Given.Land.Male, type = "diag") #


## Plot the model ####
# Standardized visit number
effect7 <- Effect(focal.predictors=c("Treatment","VisitNum"),Buzz.Given.Land.Male)
effect7 <- as.data.frame(effect7)

# Rename the figure legends
effect7$Treatment = plyr::mapvalues(effect7$Treatment, from = c("Inc_Benefit", "Inc_Cost", "Inc_Both", "Mod_Control"), 
                                    to = c("Benefit", "Cost","Both","Control"))

# Plot model with 95% CI
Figure2b <- ggplot(effect7, aes(x=VisitNum, y=fit, color=Treatment, fill=Treatment)) + 
  geom_line(lwd=1) +
  geom_ribbon(aes(ymin=lower, ymax=upper), alpha=0.2, colour=NA) + 
  scale_linetype_manual(values=c("solid","solid","solid")) + 
  ylab("Proportion lands with buzzing") + xlab("Visit number") + theme_classic() + 
  theme(text=element_text(size=14)) +
  scale_color_viridis_d(name = "Treatment", end = 0.7, option = "inferno") + 
  scale_fill_viridis_d(name = "Treatment", end =0.7, option = 'inferno') +
  theme(legend.position  =c(0.5, 0.74), legend.direction = "horizontal", 
        legend.background = element_rect(fill="transparent", colour = NA),
        strip.background = element_blank(),
        legend.title = element_text(size = 10), legend.text = element_text(size = 10),
        panel.spacing.x = unit(1, "lines"), 
        plot.margin = margin(c(0.2,0.51, 0.2, 0.2), unit = "cm")) + 
  guides(colour=guide_legend(nrow=2, byrow=TRUE)) +
  xlim(0,60) + ylim(0,1)
plot(Figure2b) #To see the figure in R

# Set date
DATE <- "5-5-2023"

# Figure dimensions and name
tiff(filename=paste("Fig2b_BuzzGivenLands_Male_60vis_", DATE, ".tif", sep=""), units="mm", width=85, height=85, res=1200)

# Export the figure
plot(Figure2b)

# Close the plotting device
setwd(PLOTDIR); 
dev.off()
#_______________________________________________________________________________________________________________________________#


#_______________________________________________________________________________________________________________________________#
# QUESTION 6 ####
# Do bees BUZZ on females more frequently with experience and does this depend on different adjustments of costs and benefits?
# Analyze only FEMALES. 

# Run the model
# Nested random effects; NumVisits within Bee within Colony
Buzz.Given.Land.Female <- glmmTMB(BuzzGivenLand ~ Treatment * VisitNum + (1 | ColonyID / BeeID / VisitNum), family = binomial, data = dataset6)

# Examine the model
summary(Buzz.Given.Land.Female)

# Check Type II sums of squares, for Wald Tests
Anova(Buzz.Given.Land.Female)

#Response: BuzzGivenLand
#                      Chisq Df Pr(>Chisq)    
# Treatment          43.555  3  1.876e-09 ***
# VisitNum           20.067  1  7.479e-06 ***
# Treatment:VisitNum 31.286  3  7.400e-07 ***

# Post-hoc test
emmeans(Buzz.Given.Land.Female, list(pairwise ~ Treatment), adjust = "tukey")

# $`pairwise differences of Treatment`
# 1                         estimate    SE   df t.ratio p.value
# Inc_Benefit - Inc_Both       1.984 0.454 1169   4.371  0.0001
# Inc_Benefit - Inc_Cost       1.728 0.492 1169   3.511  0.0026
# Inc_Benefit - Mod_Control   -0.386 0.415 1169  -0.930  0.7889
# Inc_Both - Inc_Cost         -0.256 0.496 1169  -0.517  0.9551
# Inc_Both - Mod_Control      -2.370 0.456 1169  -5.196  <.0001
# Inc_Cost - Mod_Control      -2.113 0.493 1169  -4.285  0.0001

# Model assumptions via DHARMa
simulationOutput1 <- simulateResiduals(fittedModel = Buzz.Given.Land.Female, n = 1000)
plot(simulationOutput1) # QQ plot is good
testUniformity(simulationOutput = simulationOutput1) # Uniformity good
testDispersion(simulationOutput1) # Dispersion good
testZeroInflation(simulationOutput1) #Zero inflation good

# Model assumptions via sjPlot
# Normality of residuals and outliers
plot_model(Buzz.Given.Land.Female, type = "diag") #


## Plot the model ####
# Standardized visit number
effect6 <- Effect(focal.predictors=c("Treatment","VisitNum"),Buzz.Given.Land.Female) #Full: BIIIIIIG error bars lol
effect6 <- as.data.frame(effect6)

#Rename the figure legends
effect6$Treatment = plyr::mapvalues(effect6$Treatment, from = c("Inc_Benefit", "Inc_Cost", "Inc_Both", "Mod_Control"), 
                                    to = c("Benefit", "Cost","Both","Control"))

# Plot model with 95% CI
Figure2c <- ggplot(effect6, aes(x=VisitNum, y=fit, color=Treatment, fill=Treatment)) + 
  geom_line(lwd=1) +
  geom_ribbon(aes(ymin=lower, ymax=upper), alpha=0.2, colour=NA) + 
  scale_linetype_manual(values=c("solid","solid","solid")) + 
  ylab("Proportion lands with buzzing") + xlab("Visit number") + theme_classic() + 
  theme(text=element_text(size=14)) +
  scale_color_viridis_d(name = "Treatment", end = 0.7, option = "inferno") + 
  scale_fill_viridis_d(name = "Treatment", end =0.7, option = 'inferno') +
  theme(legend.position  =c(0.5, 0.9), legend.direction = "horizontal", 
        legend.background = element_rect(fill="transparent", colour = NA),
        strip.background = element_blank(),
        legend.title = element_text(size = 10), legend.text = element_text(size = 10),
        panel.spacing.x = unit(1, "lines"), 
        plot.margin = margin(c(0.2,0.51, 0.2, 0.2), unit = "cm")) +
        guides(colour=guide_legend(nrow=2, byrow=TRUE)) +
  xlim(0,60) + ylim(0,1) 
plot(Figure2c) #To see the figure in R

# Set date
DATE <- "5-5-2023"

# Figure dimensions and name
tiff(filename=paste("Fig2c_BuzzGivenLands_Female_60vis_", DATE, ".tif", sep=""), units="mm", width=85, height=85, res=1200)

# Export the figure
plot(Figure2c)

# Close the plotting device
setwd(PLOTDIR); 
dev.off()

#_______________________________________________________________________________________________________________________________#


#_______________________________________________________________________________________________________________________________#
# QUESTION 7 ####
# Do bees learn differently in modified control vs unmodified control
# Did bee learn to make CORRECT DECISIONS? Analyze all visits to male and female together.

# Run the model
# Nested random effects; NumVisits within Bee within Colony
All.Correct.Control <- glmmTMB(Correct ~ Treatment * VisitNum + (1 | ColonyID / BeeID / VisitNum), family = binomial, data = dataset7)

# Examine the model
summary(All.Correct.Control)

# Check Type II sums of squares, for Wald Tests
Anova(All.Correct.Control)

# Response: Correct
#                        Chisq Df Pr(>Chisq)    
# Treatment          11.5467  1  0.0006787 ***
# VisitNum            3.9563  1  0.0466968 *  
# Treatment:VisitNum  0.1296  1  0.7188416

# Model assumptions via DHARMa
simulationOutput1 <- simulateResiduals(fittedModel = All.Correct.Control, n = 1000)
plot(simulationOutput1) #Plot of QQ plot looks good
testUniformity(simulationOutput = simulationOutput1) # Uniformity good
testDispersion(simulationOutput1) # Dispersion good
testZeroInflation(simulationOutput1) #Zero inflation good

# Model assumptions via sjPlot
# Normality of residuals and outliers
plot_model(All.Correct.Control, type = "diag") #Looks normal

## Plot the model ####
# Standardized visit number
effect7 <- Effect(focal.predictors=c("Treatment","VisitNum"),All.Correct.Control)
effect7 <- as.data.frame(effect7)

#Rename the figure legends
effect7$Treatment = plyr::mapvalues(effect7$Treatment, from = c("Mod_Control", "Unmod_Control"), 
                                    to = c("Modified control", "Unmodified control"))

# Plot model with 95% CI
Figure3a <- ggplot(effect7, aes(x=VisitNum, y=fit, color=Treatment, fill=Treatment)) + 
  geom_line(lwd=1) +
  geom_ribbon(aes(ymin=lower, ymax=upper), alpha=0.2, colour=NA) + 
  scale_linetype_manual(values=c("solid","solid","solid")) + 
  ylab("Proportion correct decisions") + xlab("Visit number") + theme_classic() + 
  theme(text=element_text(size=14)) +
  scale_color_viridis_d(name = "Treatment", end = 0.7, option = "inferno") + 
  scale_fill_viridis_d(name = "Treatment", end =0.7, option = 'inferno') +
  theme(legend.position  =c(0.5, 0.9), legend.direction = "horizontal", 
        legend.background = element_rect(fill="transparent", colour = NA),
        strip.background = element_blank(),
        legend.title = element_text(size = 10), legend.text = element_text(size = 10),
        panel.spacing.x = unit(1, "lines"), 
        plot.margin = margin(c(0.2,0.51, 0.2, 0.2), unit = "cm")) + 
  guides(colour=guide_legend(nrow=2, byrow=TRUE)) +
  xlim(0,60) + ylim(0,1)
plot(Figure3a) #To see the figure in R

# Set date
DATE <- "5-5-2023"

# Figure dimensions and name
tiff(filename=paste("Fig3a_CNTRL_CorrectDecisions_60vis_", DATE, ".tif", sep=""), units="mm", width=85, height=85, res=1200)

# Export the figure
plot(Figure3a)

# Close the plotting device
setwd(PLOTDIR); 
dev.off()

#_______________________________________________________________________________________________________________________________#


#_______________________________________________________________________________________________________________________________#
# QUESTION 8 ####
# Do bees learn differently in modified control vs unmodified control
# Did bee learn to make CORRECT DETECTIONS? Analyze only visits to male.

# Run the model
# Nested random effects; VisitNum within Bee within Colony
Male.Correct.Control <- glmmTMB(Correct ~ Treatment * VisitNum + (1 | ColonyID / BeeID / VisitNum), family = binomial, data = dataset8)

# Examine the model
summary(Male.Correct.Control)

# Check Type II sums of squares, for Wald Tests
Anova(Male.Correct.Control)

# Response: Correct
#                      Chisq Df Pr(>Chisq)    
# Treatment           2.5694  1     0.1090    
# VisitNum           18.8030  1  1.449e-05 ***
# Treatment:VisitNum  3.5457  1     0.0597 .

# Model assumptions via DHARMa
simulationOutput1 <- simulateResiduals(fittedModel = Male.Correct.Control, n = 1000)
plot(simulationOutput1) # QQ plot has a significant deviation 0.01
testUniformity(simulationOutput = simulationOutput1) # Uniformity good
testDispersion(simulationOutput1) # Dispersion good
testZeroInflation(simulationOutput1) #Zero inflation good

# Model assumptions via sjPlot
# Normality of residuals and outliers
plot_model(Male.Correct.Control, type = "diag") #

## Plot the model ####
# Standardized visit number
effect8 <- Effect(focal.predictors=c("Treatment","VisitNum"),Male.Correct.Control)
effect8 <- as.data.frame(effect8)

#Rename the figure legends
effect8$Treatment = plyr::mapvalues(effect8$Treatment, from = c("Mod_Control", "Unmod_Control"), 
                                    to = c("Modified control", "Unmodified control"))


# Plot model with 95% CI
Figure3b <- ggplot(effect8, aes(x=VisitNum, y=fit, color=Treatment, fill=Treatment)) + 
  geom_line(lwd=1) +
  geom_ribbon(aes(ymin=lower, ymax=upper), alpha=0.2, colour=NA) + 
  scale_linetype_manual(values=c("solid","solid","solid")) + 
  ylab("Proportion correct detections") + xlab("Visit number") + theme_classic() + 
  theme(text=element_text(size=14)) +
  scale_color_viridis_d(name = "Treatment", end = 0.7, option = "inferno") + 
  scale_fill_viridis_d(name = "Treatment", end =0.7, option = 'inferno') +
  theme(legend.position  =c(0.5, 0.15), legend.direction = "horizontal", 
        legend.background = element_rect(fill="transparent", colour = NA),
        strip.background = element_blank(),
        legend.title = element_text(size = 10), legend.text = element_text(size = 10),
        panel.spacing.x = unit(1, "lines"), 
        plot.margin = margin(c(0.2,0.51, 0.2, 0.2), unit = "cm")) + 
  guides(colour=guide_legend(nrow=2, byrow=TRUE)) +
  xlim(0,60) + ylim(0,1)
plot(Figure3b) #To see the figure in R

# Set date
DATE <- "5-5-2023"

# Figure dimensions and name
tiff(filename=paste("Fig3b_CNTRL_CorrectDetections_60vis_", DATE, ".tif", sep=""), units="mm", width=85, height=85, res=1200)

# Export the figure
plot(Figure3b)

# Close the plotting device
setwd(PLOTDIR); 
dev.off()

#_______________________________________________________________________________________________________________________________#


#_______________________________________________________________________________________________________________________________#
# QUESTION 9 ####
# Do bees learn differently in modified control vs unmodified control?
# Did bee learn to make CORRECT REJECTIONS? Analyze only visits to female. 
# Run the model


# Nested random effects; NumVisits within Bee within Colony
Female.Correct.Control <- glmmTMB(Correct ~ Treatment * VisitNum + (1 | ColonyID / BeeID / VisitNum), family = binomial, data = dataset9)

# Examine the model
summary(Female.Correct.Control)

# Check Type II sums of squares, for Wald Tests
Anova(Female.Correct.Control)

#Response: Correct
#                       Chisq Df Pr(>Chisq)    
# Treatment           6.6611  1   0.009854 ** 
# VisitNum           43.0480  1  5.341e-11 ***
# Treatment:VisitNum  2.5770  1   0.108424

# Model assumptions via DHARMa
simulationOutput1 <- simulateResiduals(fittedModel = Female.Correct.Control, n = 1000)
plot(simulationOutput1) # QQ plot is good
testUniformity(simulationOutput = simulationOutput1) # Uniformity good
testDispersion(simulationOutput1) # Dispersion good
testZeroInflation(simulationOutput1) #Zero inflation good

# Model assumptions via sjPlot
# Normality of residuals and outliers
plot_model(Female.Correct.Control, type = "diag") #

## Plot the model ####
# Standardized visit number
effect9 <- Effect(focal.predictors=c("Treatment","VisitNum"),Female.Correct.Control)
effect9 <- as.data.frame(effect9)

#Rename the figure legends
effect9$Treatment = plyr::mapvalues(effect9$Treatment, from = c("Mod_Control", "Unmod_Control"), 
                                    to = c("Modified control", "Unmodified control"))


# Plot model with 95% CI
Figure3c <- ggplot(effect9, aes(x=VisitNum, y=fit, color=Treatment, fill=Treatment)) + 
  geom_line(lwd=1) +
  geom_ribbon(aes(ymin=lower, ymax=upper), alpha=0.2, colour=NA) + 
  scale_linetype_manual(values=c("solid","solid","solid")) + 
  ylab("Proportion correct rejections") + xlab("Visit number") + theme_classic() + 
  theme(text=element_text(size=14)) +
  scale_color_viridis_d(name = "Treatment", end = 0.7, option = "inferno") + 
  scale_fill_viridis_d(name = "Treatment", end =0.7, option = 'inferno') +
  theme(legend.position  =c(0.5, 0.9), legend.direction = "horizontal", 
        legend.background = element_rect(fill="transparent", colour = NA),
        strip.background = element_blank(),
        legend.title = element_text(size = 10), legend.text = element_text(size = 10),
        panel.spacing.x = unit(1, "lines"), 
        plot.margin = margin(c(0.2,0.51, 0.2, 0.2), unit = "cm")) + 
  guides(colour=guide_legend(nrow=2, byrow=TRUE)) +
  xlim(0,60) + ylim(0,1)
plot(Figure3c) #To see the figure in R

# Set date
DATE <- "5-5-2023"

# Figure dimensions and name
tiff(filename=paste("Fig3c_CNTRL_CorrectRejections_60vis_", DATE, ".tif", sep=""), units="mm", width=85, height=85, res=1200)

# Export the figure
plot(Figure3c)

# Close the plotting device
setwd(PLOTDIR); 
dev.off()
#_______________________________________________________________________________________________________________________________#


#_______________________________________________________________________________________________________________________________#
# QUESTION 10 ####
# Do bees learn differently in modified control vs unmodified control?
# Analyze all LANDS, proportion that are on males vs females.

# Run the model
# Nested random effects; NumVisits within Bee within Colony
All.Male.Land.Control <- glmmTMB(LandsOnMale ~ Treatment * VisitNum + (1 | ColonyID / BeeID / VisitNum), family = binomial, data = dataset10)

# Examine the model
summary(All.Male.Land.Control)

# Check Type II sums of squares, for Wald Tests
Anova(All.Male.Land.Control)

# Response: LandsOnMale
#                      Chisq Df Pr(>Chisq)  
# Treatment          5.8249  1     0.0158 *
# VisitNum           0.6018  1     0.4379  
# Treatment:VisitNum 0.0439  1     0.8340

# Model assumptions via DHARMa
simulationOutput1 <- simulateResiduals(fittedModel = All.Male.Land.Control, n = 1000)
plot(simulationOutput1) # QQ plot is good
testUniformity(simulationOutput = simulationOutput1) # Uniformity good
testDispersion(simulationOutput1) # Dispersion good
testZeroInflation(simulationOutput1) #Zero inflation good

# Model assumptions via sjPlot
# Normality of residuals and outliers
plot_model(All.Male.Land.Control, type = "diag") #


## Plot the model ####
# Standardized visit number
effect10 <- Effect(focal.predictors=c("Treatment","VisitNum"),All.Male.Land.Control)
effect10 <- as.data.frame(effect10)

#Rename the figure legends
effect10$Treatment = plyr::mapvalues(effect10$Treatment, from = c("Mod_Control", "Unmod_Control"), 
                                     to = c("Modified control", "Unmodified control"))


# Plot model with 95% CI
Figure3d <- ggplot(effect10, aes(x=VisitNum, y=fit, color=Treatment, fill=Treatment)) + 
  geom_line(lwd=1) +
  geom_ribbon(aes(ymin=lower, ymax=upper), alpha=0.2, colour=NA) + 
  scale_linetype_manual(values=c("solid","solid","solid")) + 
  ylab("Proportion lands on models") + xlab("Visit number") + theme_classic() + 
  theme(text=element_text(size=14)) +
  scale_color_viridis_d(name = "Treatment", end = 0.7, option = "inferno") + 
  scale_fill_viridis_d(name = "Treatment", end =0.7, option = 'inferno') +
  theme(legend.position  =c(0.5, 0.9), legend.direction = "horizontal", 
        legend.background = element_rect(fill="transparent", colour = NA),
        strip.background = element_blank(),
        legend.title = element_text(size = 10), legend.text = element_text(size = 10),
        panel.spacing.x = unit(1, "lines"), 
        plot.margin = margin(c(0.2,0.51, 0.2, 0.2), unit = "cm")) + 
  guides(colour=guide_legend(nrow=2, byrow=TRUE)) +
  xlim(0,60) + ylim(0,1)
plot(Figure3d) #To see the figure in R

# Set date
DATE <- "5-5-2023"

# Figure dimensions and name
tiff(filename=paste("Fig3d_CNTRL_ProportionLandsOnMale_60vis_", DATE, ".tif", sep=""), units="mm", width=85, height=85, res=1200)

# Export the figure
plot(Figure3d)

# Close the plotting device
setwd(PLOTDIR); 
dev.off()
#_______________________________________________________________________________________________________________________________#


#_______________________________________________________________________________________________________________________________#
# QUESTION 11 ####
# Does quinine or pollen supplementation affect what flower naive bees prefer on their first visit?
# Analyze first landing for each treatment, male vs female

# G-tests
# Run the models

Inc_Benefit = c(7,12) #Sums to the total N
GTest(Inc_Benefit) 
# G = 1.3314, X-squared df = 1, p-value = 0.2486

Inc_Both = c(11,9) #Sums to the total N
GTest(Inc_Both) 
# G = 0.20033, X-squared df = 1, p-value = 0.6545

Inc_Cost = c(10,10) #Sums to the total N
GTest(Inc_Cost) 
# G = 0, X-squared df = 1, p-value = 1

Mod_Control = c(12,9) #Sums to the total N
GTest(Mod_Control) 
# G = 0.43004, X-squared df = 1, p-value = 0.512

Unmod_Control = c(5,11) #Sums to the total N
GTest(Unmod_Control) 
# G = 2.3059, X-squared df = 1, p-value = 0.1289
```
